# COUSIN (COdon Usage Similarity INdex): A normalized measure of Codon Usage Preferences

**DOI:** 10.1101/600361

**Authors:** Jérôme Bourret, Samuel Alizon, Ignacio G. Bravo

## Abstract

Codon Usage Preferences (CUPrefs) describe the unequal usage of synonymous codons at the gene, genomic region or genome scale. Numerous indices have been developed to measure the CUPrefs of a sequence. We introduce a normalized index to calculate CUPrefs called COUSIN for COdon Usage Similarity INdex. This index compares the CUPrefs of a query against those of a reference dataset and normalizes the output over a Null Hypothesis of random codon usage. COUSIN results can be easily interpreted, quantitatively and qualitatively. We exemplify the use of COUSIN and highlight its advantages with an analysis on the complete coding sequences of eight divergent genomes, two of them with extreme nucleotide composition. Strikingly, COUSIN captures a hitherto unreported bimodal distribution in CUPrefs in genes in the human and in the chicken genomes. We show that this bimodality can be explained by the global nucleotide composition bias of the chromosome in which the gene resides, and by the precise location within the chromosome. Our results highlight the power of the COUSIN index and uncover unexpected characteristics of the CUPrefs in human and chicken. An eponymous tool written in python3 to calculate COUSIN is available for online or local use.

## 1 Introduction

Translation of messenger RNAs (mRNA) into proteins is a central molecular biology process common to all forms of life. During translation, ribosomes proceed along the mRNA in steps of three nucleotides, called codons. While “reading” the mRNA, the ribosome allows pairing of a mRNA codon against the complementary nucleotide triplet on a transfer RNA (tRNA), catalysing the polymerisation of amino acids to synthesise peptides and proteins (Quax *et al.*, 2015). 64 nucleotide triplets are available and, in the standard genetic code, 61 codons encode for the 20 standard amino acids (Belalov and Lukashev, 2013). Because of this asymmetry, certain groups of two, three, four or six codons encode for the same amino acid. Such groups of codons are known as “synonymous codons”, and the existence of multiple coding alternatives for a single amino acid are often referred to as the degeneracy of the genetic code (Nirenberg and Matthaei, 1961; Khorana *et al.*, 1966).

Synonymous codons are not used with similar frequencies. This deviation from random use is known as Codon Usage Preferences (CUPrefs) or Codon Usage Bias (CUB). Deviations from random usage of synonymous codons occur between nucleotide stretches within a gene, between genes within a genome and between genomes in different organisms (Grantham *et al.*, 1980; Carbone *et al.*, 2003). Since codons are the units of information integration during translation, it was originally proposed for *Escherichia coli* that a connection may exist between CUPrefs and the overall efficiency of the translation process (Gouy and Gautier, 1982; Sharp and Li, 1987). Under this assumption, the presence of a given synonymous codon at a given location could be explained by natural selection (Grantham *et al.*, 1980; Bennetzen and Hall, 1982; Sharp and Li, 1987). In rapidly growing unicellular organisms, variations in the tRNA pools have been hypothesised to fuel such evolutionary forces, supported by the fact that in these organisms the CUPrefs match well tRNA abundance in the cell (Ikemura, 1981; Akashi, 1994). Additionally, mutational bias shapes CUPrefs by modifying nucleotide frequencies and thereby codon frequencies (Knight *et al.*, 2001; Urrutia and Hurst, 2001; Roth *et al.*, 2012), while GC-biased gene conversion leads to regional CUPrefs bias by promoting asymmetric GC-rich chromosome fragment replacement during meiotic recombination (Pouyet *et al.*, 2017; Galtier *et al.*, 2018). For example, in chromosome stretches with strong nucleotide composition bias, known as isochores in Vertebrates, differences in GC content may be the main driver of CUPrefs (Costantini *et al.*, 2006; Roth *et al.*, 2012).

A variety of indices have been developed since the 1980s to evaluate the CUPrefs of a sequence (Ikemura, 1981; Freire-Picos *et al.*, 1994; Urrutia and Hurst, 2001). Most of them compare the CUPrefs of a query against a reference set or against a Null Hypothesis chosen by the user (Shields *et al.*, 1988; Lee *et al.*, 2010). New indices are still developed (Zhang *et al.*, 2012) but the “Codon Adaptation Index” (CAI) (Sharp and Li, 1987) and the “Effective Number of Codons” (ENC) (Wright, 1990) remain the most popular ones and are still being improved (Lee *et al.*, 2010; Satapathy *et al.*, 2017). Problematically, most CUPrefs indices have little reliability when analyzing sequences with either short length, strong GC content or strong amino acid composition bias (Roth *et al.*, 2012). Furthermore, certain CUPrefs scores have limited biological meaning, and often require a certain knowledge of the studied organism to be interpreted correctly. For example, the FOP index requires the specification of a set of optimal codons (*e.g.* by determining the gene copy number of each tRNA in the studied organism) (Ikemura, 1981).

Concomitantly to the development of new CUPrefs indices, numerous software packages to evaluate CUPrefs have been implemented, such as `INCA` (Supek and Vlahovicek, 2004), `JCAT` (Grote *et al.*, 2005) and `CodonW` (Peden and Sharp, 2005). Even if most of these packages only compute the CAI and sometimes the ENC indices, some feature new and exclusive methods such as CodonO and the “Synonymous Codon Usage Order” (SCUO) score (Wan *et al.*, 2004; Angellotti *et al.*, 2007). Still, a number of indices, such as the scaled *χ*^2^ (Shields *et al.*, 1988) or the “Maximum-likelihood Codon Bias” (MCB) (Urrutia and Hurst, 2001), have never been made available to the scientific community via a dedicated software. To date, CodonW is the most complete software but it only displays outputs related to four CUPrefs indices (Peden and Sharp, 2005). This illustrates the need for a software capable of calculating CUPrefs for a wide set of indices. A final feature lacking in most softwares is the ability to perform statistical analyses, such as those developed in the `e-cai` server to assess the significance of CUPrefs differences between a query and a reference dataset (Puigbï¿œ *et al.*, 2008b).

We introduce here `COUSIN` (acronym for COdon Usage Similarity INdex), a novel index conceived to estimate CUPrefs with a straightforward biological interpretation. We implement this index together with seven other existing ones in an eponym Python3 software that is available for local or online use. To illustrate all the potentialities of `COUSIN`, we compare it to the well known CAI when analyzing eight complete Coding DNA Sequence (CDSs) datasets from a range of organisms with large differences in nucleotide composition and genome organization.

## 2 New Approaches

### 2.1 Measuring Codon Usage Preferences

In this section, we introduce two versions of our COUSIN index (COUSIN_18_ and COUSIN_59_) and present CAI18, a modification of the CAI index introduced by Sharp et li Sharp and Li (1987) to allow comparison with COUSIN_18_. The notations used to define these indexes are given in Table 1.

**Table 1:** Notations used to define COUSIN and CAI indexes

| Symbol | Description |
| --- | --- |
| $c$ | Codon |
| $a$ | Amino acid |
| $f$ | Frequency |
| ref | Reference |
| que | Query |
| $H_0$ | Null Hypothesis |
| $L$ | Query length |
| $k_a$ | Synonymous codons of the amino acid $a$ |
| $\mathcal{A}$ | Amino acids existing in both query and reference |
| $\mathcal{N}$ | The number of amino acids existing in both query and reference |

#### 2.1.1 The COUSIN index

We conceived COUSIN to evaluate the CUPrefs of a sequence while offering biologically meaningful results: the CUPrefs of a query are compared to those of a reference dataset, and the results of this comparison are normalized over a Null Hypothesis which assumes a random usage of synonymous codons.

The COUSIN score calculation involves four steps:

1. Calculate deviation scores (dev_*c,a*_) for each codon (*c*) of each amino acid (*a*) in the reference dataset, compared to the Null Hypothesis:

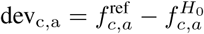

where 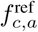 is the frequency of the codon *c* in the reference dataset and 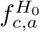 the corresponding frequency under the Null Hypothesis.
2. Define a weight for each codon (W_*c,a*_), by multiplying the codon frequency in the reference by its deviation score:

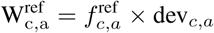
3. Repeat step 2 for the codon frequencies in the query:

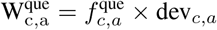

Using the same deviation score to calculate the weights allows us to compare the scores of the query and of the reference.
4. The 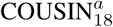 score of each amino acid is the ratio of the sum of the weights of all synonymous codons for this amino acid in the query dataset over the corresponding sum of the weights in the reference dataset:

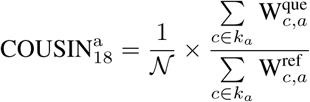

where *𝒩* is the number of amino acids present in both the query and the reference.
5. The global COUSIN score is obtained by adding the COUSIN scores of all amino acids found in both the query and the reference:

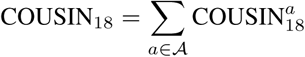

where *𝒜* is the set of amino acids present in both the query and the reference.

By design, the results of COUSIN have an immediate interpretation and are directly suitable for hypothesis testing (Figure 1). COUSIN scores can be compared against two threshold values: a COUSIN score of 1 indicates that the CUPrefs in the query are similar to those in reference dataset, while a COUSIN score of 0 indicates that the CUPrefs in the query are similar to those in the Null Hypothesis (*i. e.* random usage of synonymous codons). Other COUSIN scores outside these two values can be interpreted as follows:

**Figure 1:**
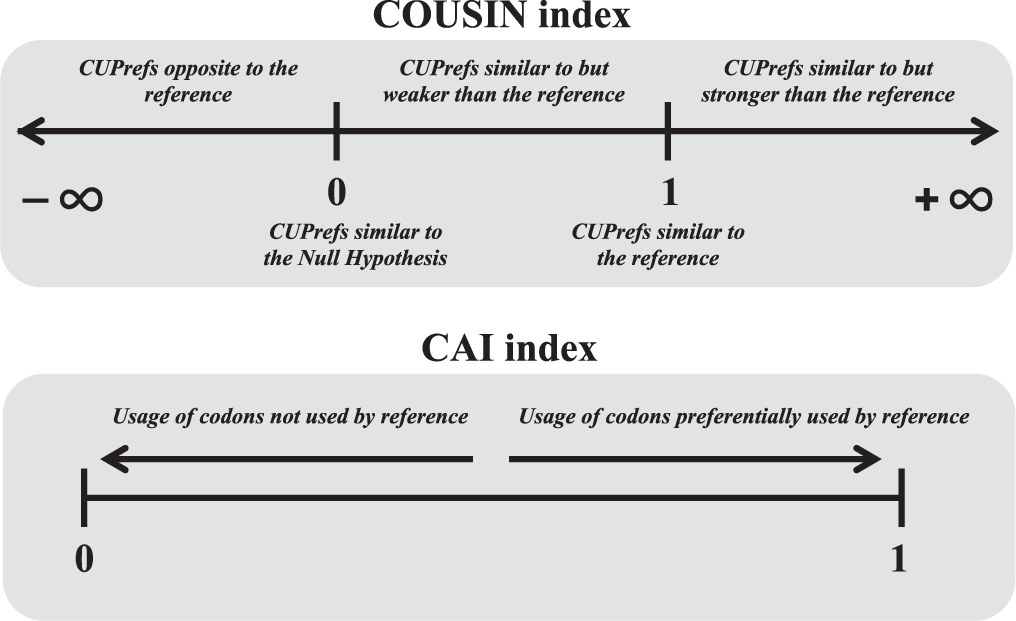
Range of values and interpretation of COUSIN (top scale) and CAI (bottom scale) indexes

- a COUSIN score above 1 indicates that CUPrefs in the query are similar to those in the reference but of larger magnitude, *i. e.* the more frequent codons in the reference are even more frequently used in the query;
- a COUSIN score between 0 and 1 indicates that CUPrefs in the query are similar to those in the reference but of smaller magnitude, *i. e.* the more frequent codons in the reference are used in the query more often than in the Null hypothesis of equal frequency;
- a COUSIN score below 0 indicates that CUPrefs in the query are opposite to those in the reference, *i. e.* the less used codons in the reference are used more often in the query than in the Null hypothesis of equal frequency;

#### 2.1.2 Accounting for amino acid composition in CUPrefs

It has been suggested that amino acid composition may affect the CAI score obtained for sequences with similar codon usage, such that the lower the amino acid diversity in a sequence, the higher the bias (Roth *et al.*, 2012). The version of COUSIN described above, namely COUSIN_18_, assigns equal contribution to all amino acids. We therefore conceived an alternative version of COUSIN named COUSIN_59_, that accounts for amino acid composition in the query, by weighting the contribution of each amino acid by its frequency in the query, as follows:

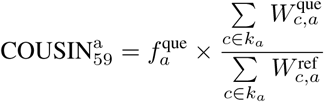

where 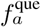 is the frequency of the amino acid *a* in the query.

The final step in the calculation of the index remains unchanged :

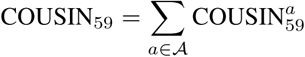

In the classical CAI score, the amino acid composition of the query sequence is included in the calculation because all codons contribute equally to the final score (see supplementary Informations 1 for a reminder of the CAI definition). This calculation is analogous to our description of COUSIN_59_, and we therefore refer to it as CAI59. For the sake of completeness, we introduce an alternative CAI definition, hereafter named CAI18, for which all amino acids contribute equally. The difference between CAI18 and CAI59 simply lies in the calculation of the geoindexal mean, as follows:

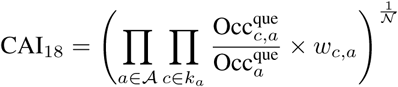

where 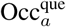 is the number of occurrences of the amino acid *a* in the query, 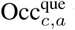 the number of occurrences of codon *c* in the query and *w*_*c,a*_ the relative adaptiveness score (Supplementary Information 1).

Both pairs COUSIN_18_ and COUSIN_59_, and CAI_18_ and CAI_59_ therefore differ in the way the amino acid composition is accounted for in the calculation. With the “18” methods, all amino acids contribute equally, independently of their frequency in the protein. These “18” methods can be envisioned as the “amino acid by amino acid” CUPrefs of a sequence. With the “59” methods, all individual codons contribute equally, so that the final contribution of each amino acid is proportional to its frequency in the protein. These “59” methods can be envisioned as the “codon by codon” CUPrefs of a sequence.

### 2.2 COUSIN software

We designed a software package to implement our new COUSIN index along with other seven existing indices to facilitate CUPrefs analysis and comparisons between methods. Importantly, our software package can also perform statistical analyses by means of sequence data simulation. The COUSIN software and its documentation are accessible online at http://COUSIN.ird.fr. A local version can be downloaded from the same website to be used on a UNIX-like Operating System. This software is coded in Python3 programming language. In its local version, it runs through a Unix terminal in the form of command lines and accepts several parameters and options. To avoid ambiguities, we will refer to the software with the notation `COUSIN`

#### 2.2.1 COUSIN architecture

The main input data for `COUSIN` are query sequences in a FASTA format. Depending on the task performed, it may be necessary to provide additional input files such as a reference dataset in a kazusa-like codon usage table format (Nakamura *et al.*, 2000). From these data, `COUSIN` performs a number of tasks either routinely or according to user specifications. At the end of a task, graphical and textual results are displayed.

Figure 2 describes the global architecture of `COUSIN`.

**Figure 2:**
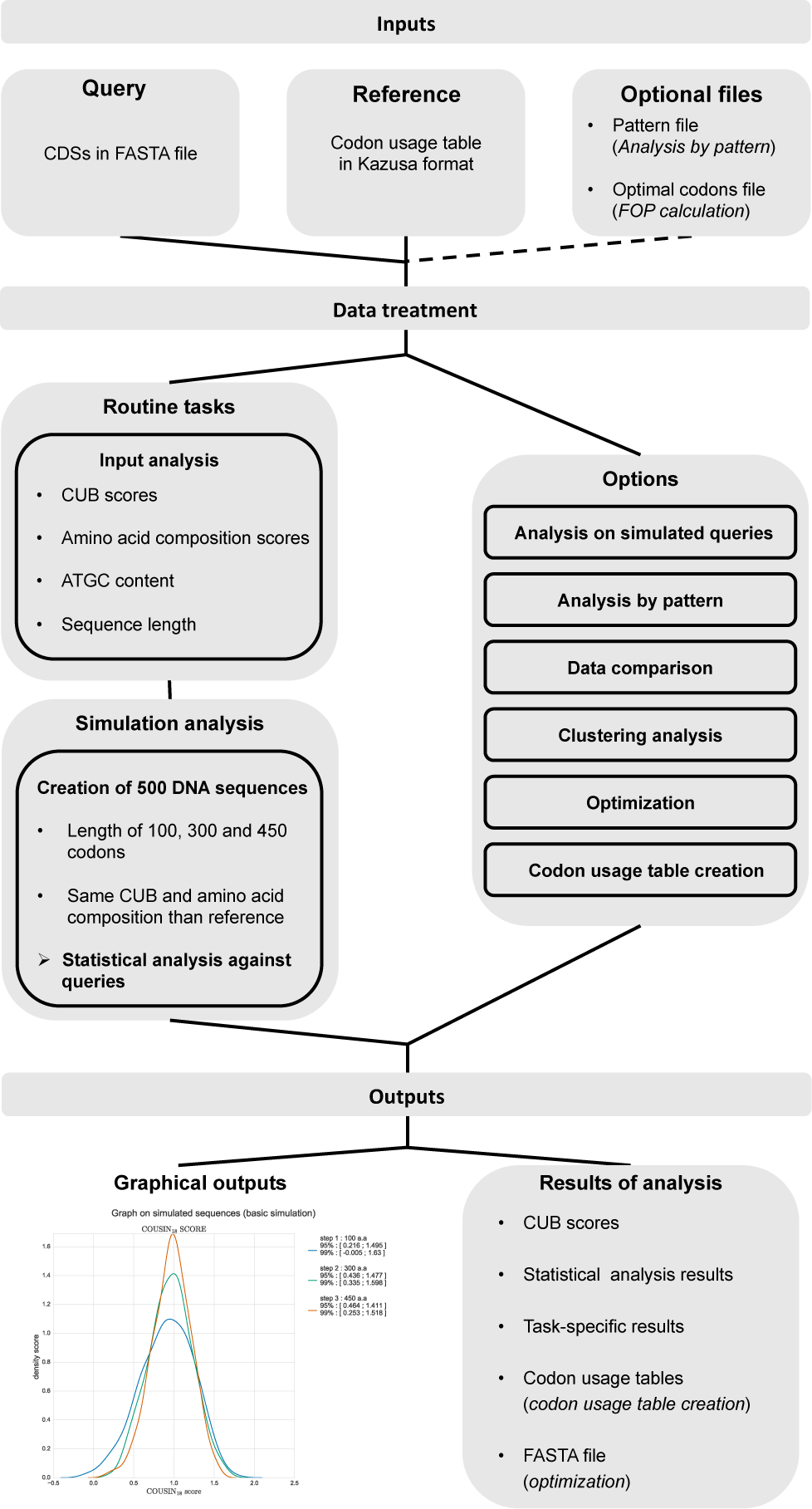
Architecture of the COUSIN software. The COUSIN software requires input data from the user such as sequences in a FASTA format and a Codon Usage Table in a kazusa-style format (Nakamura *et al.*, 2000). COUSIN then performs a CUPrefs analysis on the queries by performing routine tasks. Following user specifications, other tasks can be performed to complement the analysis. Graphical and textual outputs are given at the end of a COUSIN job.

#### 2.2.2 Available measures

`COUSIN` currently features eight indices that involve CUPrefs and two indices that involve the amino acid composition of a sequence:

- COUSIN_18_ and `COUSIN`_59_.
- CAI_18_ and CAI_59_ (Sharp and Li, 1987).
- Effective Number of Codons (ENC) (Wright, 1990).
- Synonymous Codon Usage Order (SCUO) (Angellotti *et al.*, 2007).
- Frequency of Optimal Codons (FOP) (Ikemura, 1981).
- Codon Bias Index (CBI) (Bennetzen and Hall, 1982).
- Intrinsic CoDon bias Index (ICDI) (Freire-Picos *et al.*, 1994).
- scaled *χ*^2^ (Shields *et al.*, 1988).
- GRand AVerage of HYdropathy (GRAVY), that evaluates the grand average hydropathicity of a protein (Kyte and Doolittle, 1982).
- The AROMAticity score (AROMA) that evaluates the average aromaticity of a protein (Lobry and Gautier, 1994).

#### 2.2.3 COUSIN functioning

##### Input data treatment

The first and mandatory input taken by `COUSIN` is a FASTA file containing the query sequences. The software first checks whether each sequence in the input file contains nucleotide or amino acid characters. For DNA sequences, a second check is performed to determine whether they are coding sequences: the sequence must contain a number of nucleotides that is a multiple of 3 and, if it contains a STOP codon, it must be present at the end of the sequence. If any of these conditions are not met, the sequence is removed from the analysis and the user is warned. Sequences bearing the same header than a sequence already analyzed are also put aside.

Except for the codon usage table creation and data comparison additional steps (see 2.2.3), `COUSIN` requires the user to enter a codon usage table in the kazusa-like format, to be used as reference set. `COUSIN` first validates the format of the codon usage table before verifying that it is informative. Many indices to evaluate CUPrefs perform poorly if any of the codons does not occur in the reference codon usage table, which often happens if this table is constructed from an insufficient dataset. It is therefore recommended to use a reference based on a comprehensive CDS dataset, *e.g.* the complete CDSs of the organism studied. In order to avoid comparisons against empty values, `COUSIN` replaces any null codon frequency value in the reference dataset by an approximation calculated using a non-informative prior for codon choice, as follows:

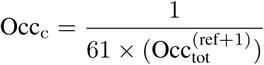

where 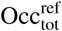 is the total number of codons found in the reference.

##### Outputs display

By default, `COUSIN` displays all scores and statistical results in a Tabulated Separated Values (TSV) format. Depending on the additional steps instructed by the user, other files can be provided. As an example, a FASTA file containing optimized sequences is given at the end of an optimization (see 2.2.3).

##### Routine tasks

For any entry `COUSIN` initially performs the following calculations:

- overall GC and nucleotide composition,
- sequence length,
- CUPrefs and amino acid composition scores for the indices described above.

If instructed by the user, `COUSIN` performs simulations to assess whether the score of a query is statistically close to that of a standard CDS encoded by the reference. To do so, it generates 500 sequences following a “random-guided” selection of amino acids and codons (Puigbï¿œ *et al.*, 2007), whereby the simulated sequences display average amino acid and CUPrefs frequencies similar to those used to construct the reference table. For each simulated sequence the CUPrefs scores are calculated and the 95% and 99% confidence intervals of the distributions of scores are estimated. The query’s score is then compared to the limits of these intervals. At the end of this step, `COUSIN` displays a graphical output that represents the range of values obtained during the simulation (Figure 3).

**Figure 3:**
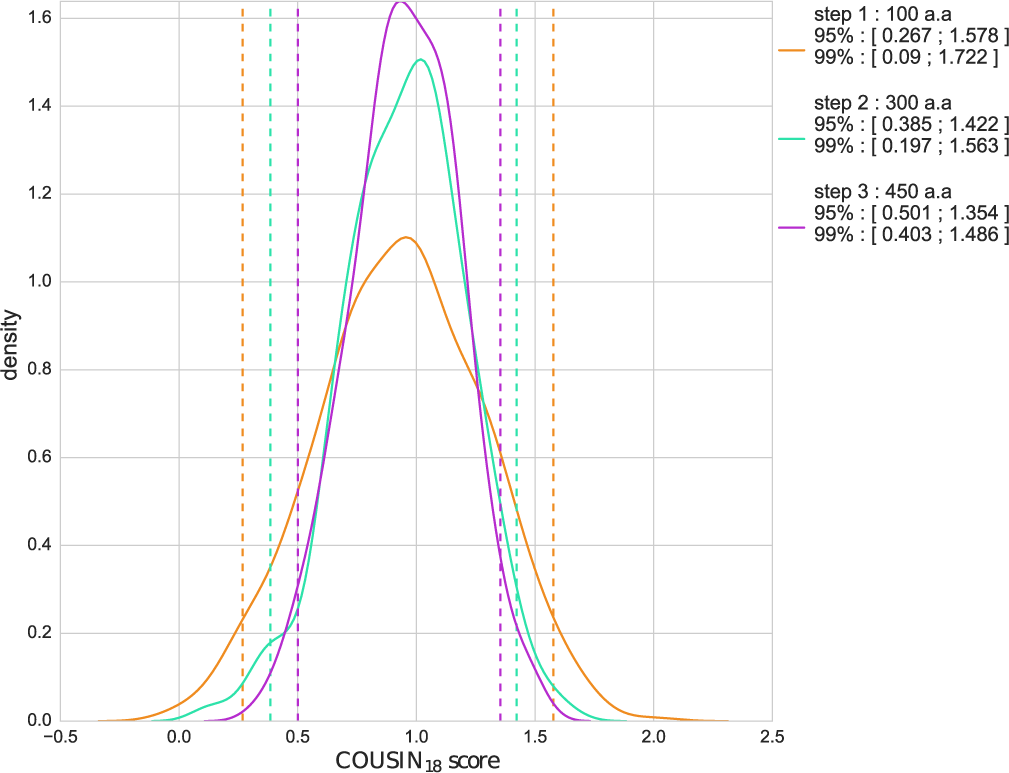
Example of the graphical output displayed by `COUSIN` software at the end of the routine steps. The density curves represent COUSIN_18_ scores of simulated sequences obtained with a “random-guided” selection following the reference CUPrefs and amino acid composition (Puigbï¿œ *et al.*, 2008a), using *E. coli* as an example. Orange, cyan and purple curves refer respectively to sequences with a number of amino acids equal to 100 (short proteins), 300 (average length of prokaryotic proteins) and 450 (average length of eukaryotic proteins). All curves are unimodal, with a mean close to 1. As expected, the longer the sequence, the lower the variance and the higher the accuracy of the scores obtained (Comeron and Aguadï¿œ, 1998; Roth *et al.*, 2012). For each curve, the legend indicates the limits of 95% and 99% confidence intervals. Dashed vertical lines indicate the respective 95% interval for orange, cyan and purple curves.

##### Additional steps

`COUSIN` proposes six additional steps to further analyse CUPrefs.

A **simulation step** related to the query. Here, two datasets are generated, each of which is built using different assumptions:

Figure 3: Example of the graphical output displayed by `COUSIN` software at the end of the routine steps. The density curves represent COUSIN_18_ scores of simulated sequences obtained with a “random-guided” selection following the reference CUPrefs and amino acid composition (Puigbï¿œ *et al.*, 2008a), using *E. coli* as an example. Orange, cyan and purple curves refer respectively to sequences with a number of amino acids equal to 100 (short proteins), 300 (average length of prokaryotic proteins) and 450 (average length of eukaryotic proteins). All curves are unimodal, with a mean close to 1. As expected, the longer the sequence, the lower the variance and the higher the accuracy of the scores obtained (Comeron and Aguadï¿œ, 1998; Roth *et al.*, 2012). For each curve, the legend indicates the limits of 95% and 99% confidence intervals. Dashed vertical lines indicate the respective 95% interval for orange, cyan and purple curves.

1. Simulated sequences having the same length as the query. In this case, the simulation follows a random-guided selection for both amino acids and codons based on the average amino acid composition and CUPRefs in the reference.
2. Simulated sequences having the same length and amino acid composition as the query. In this case, the random-guided method only selects codons that follow the CUPrefs of the reference (the amino acid composition can vary).

The distribution of CUPrefs scores of each of these two datasets is then calculated and the query’s score is compared to the 95% and 99% confidence intervals of these two distributions. If it belongs to one dataset interval and not the other, this suggests that amino acid composition significantly impacts the CUPrefs score of the query. A Wilcoxon-Mann-Whitney U-test is performed on the two simulated datasets to check whether or not they have the same median score.

An analysis of the query dataset following **header patterns** (the dataset must contain multiple queries). In this additional step, the user must submit a “pattern file” that contains a list of header patterns to `COUSIN`. Queries with the same patterns in their headers are analyzed altogether.

A **clustering step** consisting in a K-means / X-means analysis on a set of variables obtained at the end of a `COUSIN` analysis (such as CUPrefs scores, GC content, length of query or frequencies of synonymous codons) (Pelleg and Moore, 2000). The clustering results are then projected on the two first axis of a Principal Component analysis (PCA) performed in parallel. It is also possible to use a pattern file, similar to the header pattern analysis described above, to create specific clusters that are highlighted on the PCA results. In addition, a hierarchical clustering is performed. All the clustering results are stored in a file containing the header of each sequence analyzed and its corresponding cluster. Clustering graphs are also displayed at the end of the analysis.

A **sequence optimization step**, the philosophy of which is similar to that of existing software packages (Puigbï¿œ *et al.*, 2007; Grote *et al.*, 2005; Supek and Vlahovicek, 2004). This additional step modifies the CUPrefs of a query to follow those in the reference. With `COUSIN`, three different types of optimizations can be performed:

- A “Random Guided” optimization, where the codons are selected based on their frequency in the reference. This randomization can be guided towards a selection of synonymous codons maximizing GC or AT content.
- A “Random” optimization, where a codon is randomly selected among the synonymous ones for each amino acid of the sequence. This randomization can be guided towards a selection of synonymous codons maximizing GC or AT content.
- A “One amino acid, One codon” optimization, where each amino acid is represented by a unique codon (the one with the highest or lowest frequency in the reference).

The **creation of a codon usage table** from a given dataset in a kazusa-like format from a set of FASTA sequences. Indeed, although some databases contain codon usage tables, one may still need to construct one from a specific dataset (Athey *et al.*, 2017).

A **data comparison step**, where `COUSIN` calculates the Euclidean distances between the vectors of synonymous codons or amino acids frequencies among multiple datasets. These datasets can be FASTA files or Codon Usage Tables in the kazusa-like format.

With the exception of data comparison and creation of codon usage table, each additional step is performed after the routine steps described in section 2.2.3.

## 3. Material and methods

We illustrate the potential of the `COUSIN` index and compare it to the widely used CAI one by performing an analysis on the complete CDSs of eight highly unrelated organisms with contrasted GC content.

### 3.1 Datasets

For this benchmarking, we analyzed the full CDSs from two prokaryotes (*Escherichia coli, Streptomyces coelicolor*), a plant (*Arabidopsis thaliana*),a yeast (*Saccharomyces cerevisiae*), a protist (*Plasmodium falciparum*), a bird (*Gallus gallus*) and two mammals (*Homo sapiens, Mus musculus*) (Table 2). Some of these genomes were chosen because of their particularities:

**Table 2:**
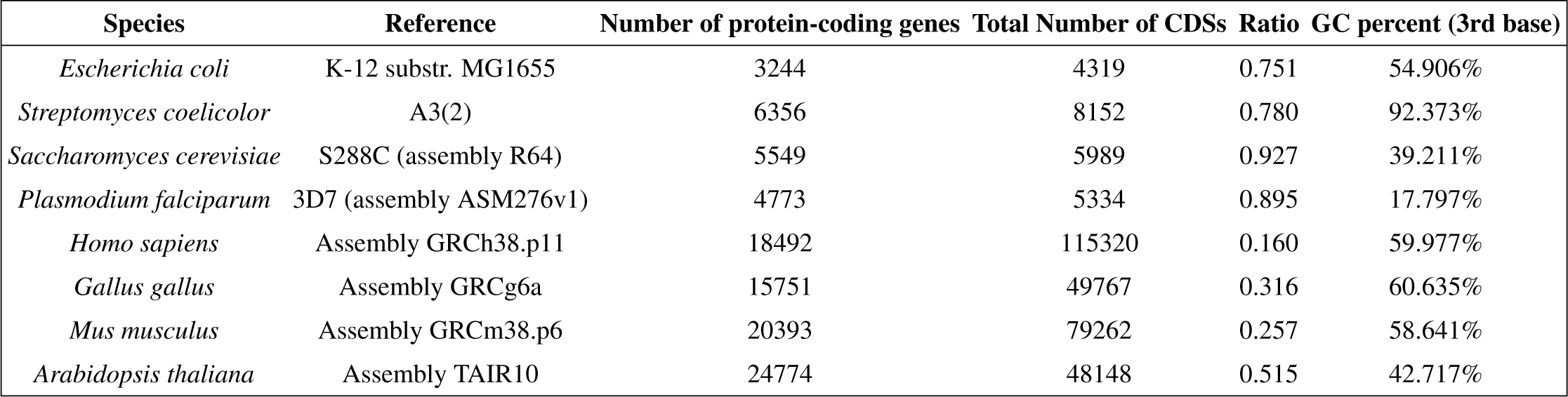
Summary statistics of the complete CDSs of the eight organisms included in the analysis. The table shows the species name, reference and accession number in the NCBI database, the number of protein-coding genes kept for the analysis (evaluated by removing isoforms and rejected sequences), the total number of CDSs retrieved (as annotated in genbank files), the ratio between the number of protein-coding genes and the total number of CDSs as well as the global GC3 content found in protein-coding genes.

- *P. falciparum* and *S. coelicolor* genomes exhibit extreme GC content;
- the *Gallus gallus* genome consists of macrochromosomes and microchrosomes (Auer *et al.*, 1987; Axelsson *et al.*, 2005);
- *Gallus gallus, Mus musculus* and *Homo sapiens* present very heterogeneous distributions of GC content within chromosomes (isochores) and/or between chromosomes (Costantini *et al.*, 2006).

### 3.2 Retrieving complete CDSs

We extracted the complete CDSs from the eight genomes using the Emboss “extractfeat” function (Rice *et al.*, 2000). For eukaryotic organisms, mitochondrial and chloroplast genomes were put aside to only keep the nuclear genome. A selection of the newly extracted CDSs was performed using the verification criteria described in section 2.2.3. To avoid redundancy during the creation of codon usage tables, only the first isoform among alternative spliced forms of a gene was kept. Finally, only CDSs with a length of at least 300 nucleotides were kept for the analyses. Indeed, most CUPrefs methods show strong biases when analyzing sequences shorter than 100 amino acids (Comeron and Aguadï¿œ, 1998; Roth *et al.*, 2012). At the end of this step, we computed the number of CDSs and the overall GC percent found at the 3rd base of each codon (GC3 content). These data are summarized in Table 2. An overview on global GC3 content of the organisms studied is given in Supplementary data 2.

### 3.3 Building reference datasets and COUSIN analysis

For each organism, we used the complete CDSs dataset to create a reference representing the average CUPrefs via the Codon Usage Table creation step proposed by `COUSIN`, and calculated the CUPrefs scores of each CDS against this reference. From this analysis, we created density curves of CAI and COUSIN scores to compare the two metrics. Finally, we performed a Pearson correlation coefficient test between COUSINand CAI scores for all CDS in each organism.

## 4. Results

### 4.1 COUSIN vs. CAI indexes

Using the Codon Usage Tables created with `COUSIN`, we calculated the COUSIN and CAI scores for each CDS of each organism. The resulting density curves are presented in Figure 4. Individual data distributions are shown in Supplementary Information 3 along with GC3 content distribution and Pearson correlation tests between this GC3 content and COUSIN_59_ or CAI_59_. The supplementary Information 4 give mean and Huber-M estimator values arising from this analysis.

**Figure 4:**
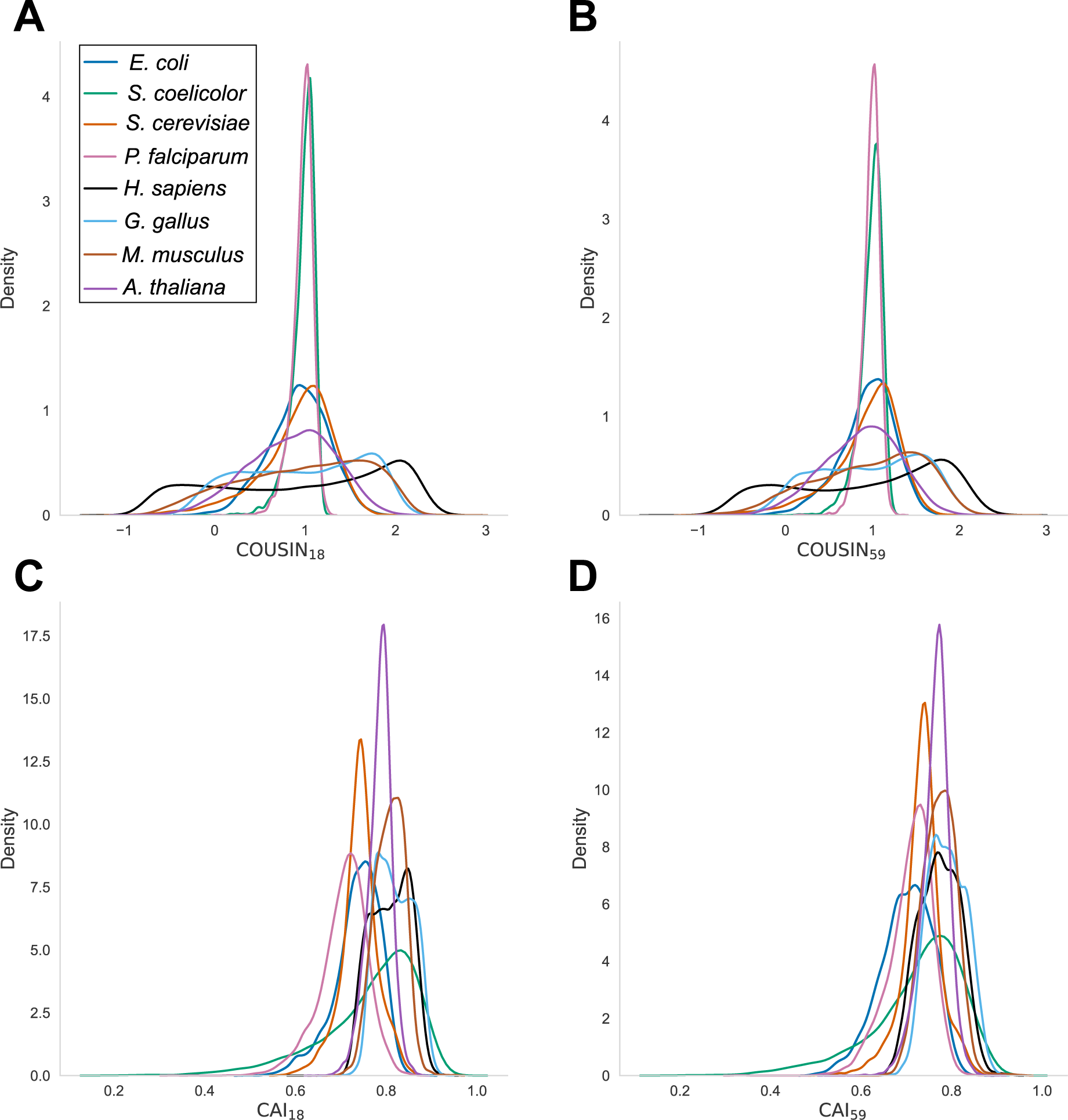
Density curves for COUSIN_59_ (A), COUSIN_59_ (B), CAI_18_ (C) and CAI59 (D) indices for the complete CDSs of the eight organisms studied (see color legend).

The analysis of CDSs with the COUSIN index highlights shared patterns as well as idiosyncrasies between organisms. All curves have a mean and Huber-M estimator score close to 1 (*i.e.* similar to that of the reference), but they strongly differ in terms of dispersion and of the global shape of data distribution, which can be unimodal (for *E. coli, S. cerevisiae, A. thaliana, S. coelicolor* and *P. falciparum*), bimodal (for *H. sapiens* and *G. gallus*) or flat with a number of local maxima (for *M. musculus)*. These differences may arise due to multiple factors such as diversity in codon usage or overall and local GC3 content. We find only few noticeable differences between COUSIN_18_ and COUSIN_59_ scores distributions, suggesting that amino acid composition has, on average, little to no impact on overall CUPrefs within each studied organism.

COUSIN values distribution curves from *S. coelicolor* and *P. falciparum* are unimodal and leptokurtic. The strong nucleotide compositional bias in these genomes (92.4% GC3 for *S. coelicolor* and 17.8% GC3 for *P. falciparum*) seems to explain these distributions with little variance, as suggested by the correlation between the two variables (Supplementary information 3, Pearson correlation scores of 0.933 for *S. coelicolor* and −0.920 for *P. falciparum*, both with p-values < 2.2^*-*16^). For other organisms with unimodal distribution but less biased nucleotide composition (*e.g. E. coli*, with 54.9% GC3), the distributions have a larger variance. The particular shapes of vertebrates curves might in turn be associated with local differences in GC3 content, as discussed below in section 4.2.

The CDSs distributions obtained with the CAI scores have unimodal shapes and exhibit differences in their mean and dispersion, with the exception of *G. gallus* and *H. sapiens* that show once again particular shapes in the distribution of scores. As for COUSIN_18_ and COUSIN_59_, CAI_18_ and CAI_59_ display few to no differences.

A major difference between COUSIN and CAI indexes resides in the immediate interpretation of the results. For COUSIN, since we compare the CUPrefs of individuals CDSs to a reference representing the overall CUPrefs of an organism, we expected an average score close to 1. For the CAI score however, in the absence of a fixed reference value, it is difficult to interpret the distribution. Moreover, the COUSIN index seems to better capture the impact of the GC3 content on CUPrefs (Supplementary Information 3). For larger genomes with strong local differences in nucleotide composition (*e.g.* chromosome isochores), the COUSIN data captures hitherto unreported patterns of CUPrefs distribution (Supplementary Informations 3).

We further compared COUSIN_59_ and CAI_59_ scores of each organism using Pearson correlation tests. The results for *E. coli* and *H. sapiens* are shown in Figure 5), and the full results are in Supplementary Information 5. For all organisms, the correlation scores between COUSIN_59_ and CAI_59_ indexes is strong and positive, ranging from 0.661 in *A. thaliana* to 0.978 in *S. coelicolor*.

**Figure 5:**
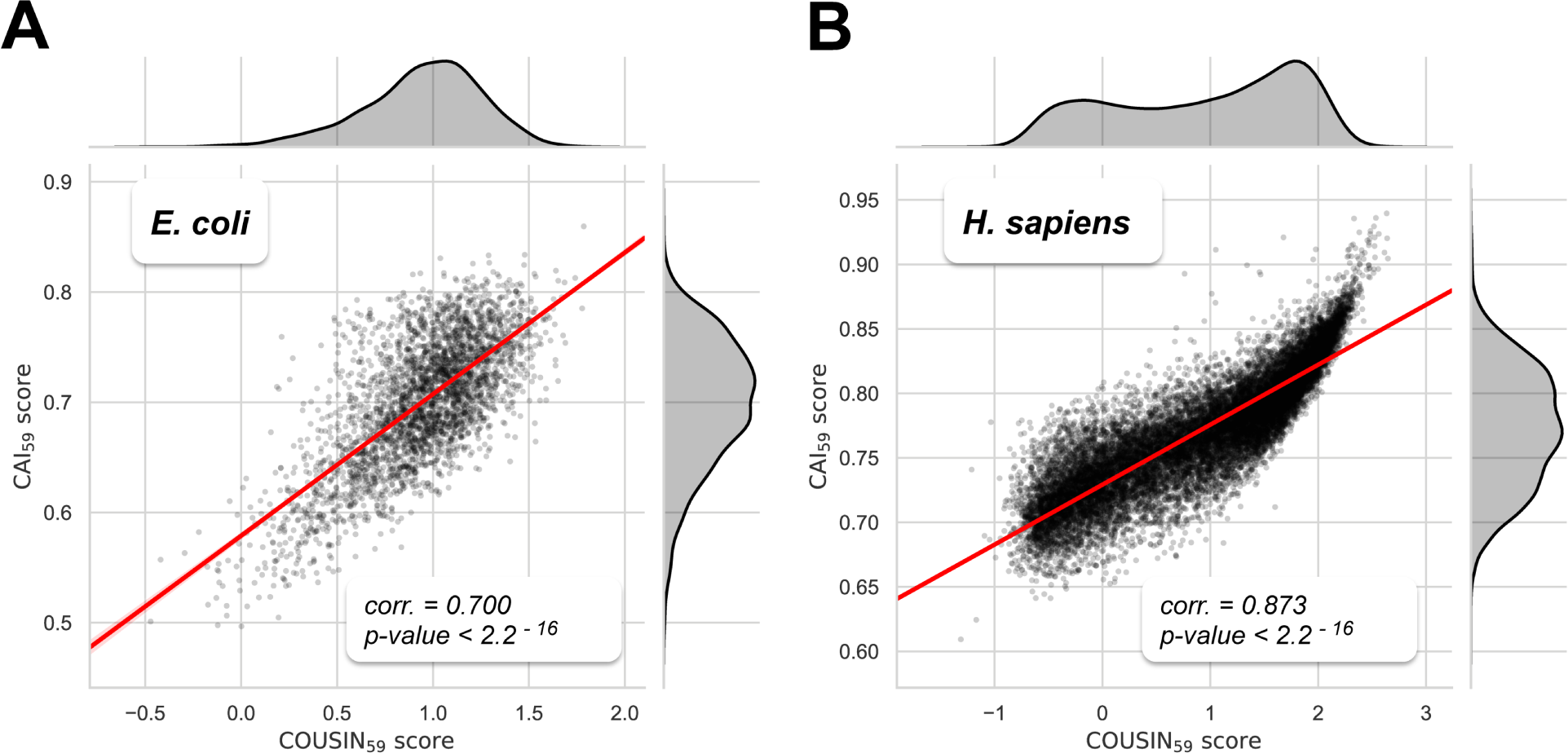
Scatter plots of *E. coli* (A) and *H. sapiens* (B) CDSs scores between COUSIN_59_ (x-axis) and CAI_59_ (y-axis) indexes. The regression line is given in red. Pearson’s correlation test results are indicated in the bottom-right of the plots. Density curves indicating the distribution of scores are shown in black at the periphery of the scatter plot.

### 4.2 COUSIN score in Vertebrates genomes

The distribution of `COUSIN`59 scores and of GC3 content in *H. sapiens, G. gallus* and *M. musculus* are strongly correlated (Supplementary Information 3 with Pearson correlation scores of 0.940, 0.818 and 0.899, all with p-values < 2.2^*-*16^). The multiple peaks observed in *H. sapiens* and *G. gallus* distributions correspond to populations of CDSs with similar GC3 content, and we hypothesise that this reflects the genomic nucleotide composition heterogeneity within or between chromosomes. Indeed, isochores in vertebrates correspond to chromosomal stretches with relativelyhomogeneous and strongly biased GC3 content (Costantini *et al.*, 2006) and microchromosomes of birds are more GC-rich than macrochromosomes (Auer *et al.*, 1987; Axelsson *et al.*, 2005). To test our hypothesis, we stratified each organism’s CDSs in three categories based on their COUSIN_59_ score:

- “Top” CDSs are the 20% ones with the highest COUSIN_59_ score.
- “Bottom” CDSs are the 20% ones with the COUSIN_59_ score.
- “Middle” CDSs are the remaining 60 % of CDSs.

Using the `KaryoploteR` package in `R`, we explored the relationship between the `COUSIN` and CAI scores, the GC3 content, the isochores regions and the position of CDSs inside *H. sapiens* chromosomes (Figure 6). As anticipated, a CDS’ GC3 content is closely related to its position in the chromosome: GC-rich CDSs are more often found in GC-rich isochores with an opposite trend for AT-rich CDSs. Furthermore, “Top” CDSs are found in GC-rich isochores but are rare in AT-rich regions. “Middle-bottom” CDSs COUSIN scores are more often found in AT-rich regions, but are still present in GC-rich isochores. This distribution is not surprising, since in *H. sapiens*, the overall CUPrefs lean towards GC-rich synonymous codons, most likely reflecting the impact of GC-biased gene conversion (Pouyet *et al.*, 2017; Galtier *et al.*, 2018). Therefore, GC-rich CDSs, which are mainly found in GC-rich regions, tend to have a higher `COUSIN` score than the CDSs found in AT-rich regions. Finally, we note that AT-rich regions contain less CDSs than GC-rich ones.

**Figure 6:**
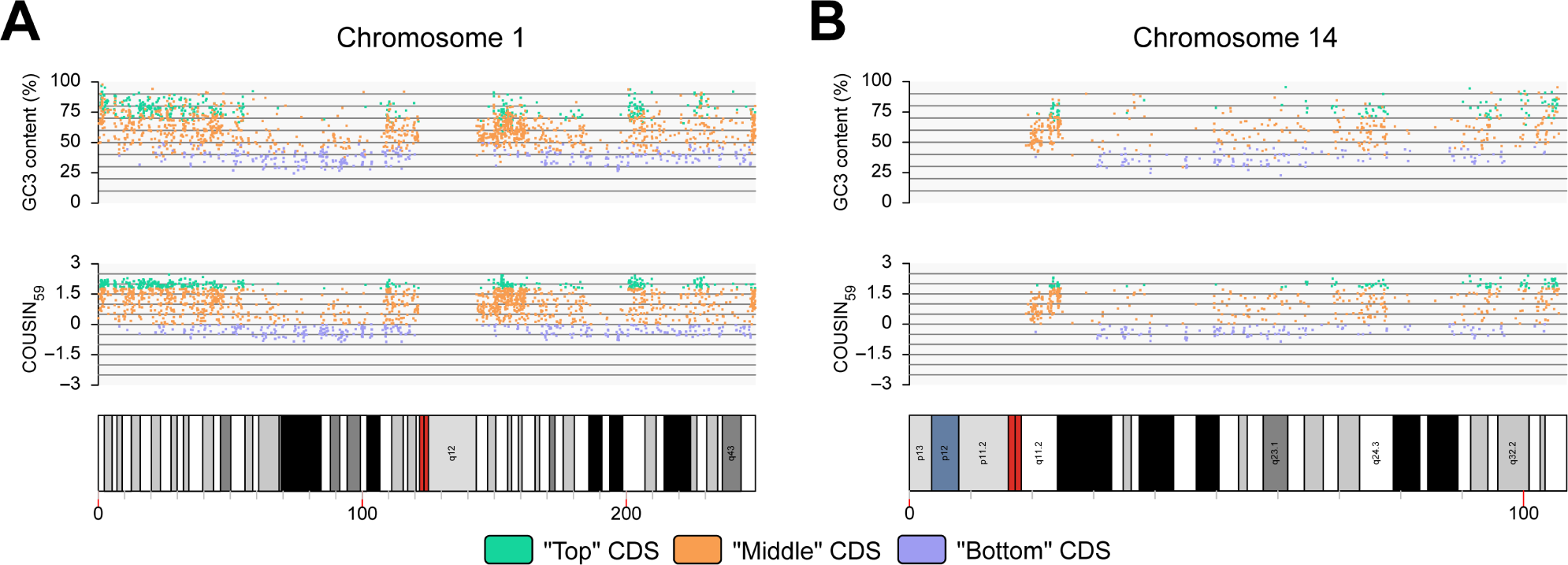
GC3 content scores (upper panel), `COUSIN`59 scores (middle panel) and structural information (lower panel) of CDSs from *H. sapiens* chromosomes 1 (A) and 14 (B). Dot colours indicate “Top” (green), “Middle” (orange) or “Bottom” (purple) CDSs. The x-axis shows the position of the CDSs in the chromosome by the distance from the p-terminal to the q-terminal in Megabases. On the lower panels, colours indicate centromeres (red), isochores (with a scale going from white for GC-rich isochores to black for AT-rich ones) and chromosomes particularities such as secondary constrictions (blue).

We further investigated changes in GC3 content and COUSIN score between chromosomes in the case of the *G. gallus* genome, which exhibits a large heterogeneity in chromosome size and a clear connection between chromosome size and both overall GC3 content and COUSIN_59_ scores (Figure 7A and B). Figure 9 shows the COUSIN_59_ and GC3 content Huber-M estimator values for each chromosome against their size (Huber *et al.*, 1981). We find a clear correlation between the size of a chromosome and both the overall GC3 content and COUSIN_59_ of the CDSs it contains: the smaller the chromosome, the higher its overall GC3 content and COUSIN_59_ score (Figure 9 A and B, Spearman correlation scores of −0.863 for GC3 content and of −0.821 for COUSIN_59_, with p-values of 5.113^*-*11^ and of 2.698^*-*9^). Similarly, the smaller the chromosome, the higher the number of “Top” CDSs and the lower the number of “Bottom” CDSs (Figure 7C). A similar but weaker trend is observed in *H. sapiens* genome, for which overall GC3 content and COUSIN_59_ score also seem to correlate negatively with chromosome size (Figure 9C and D, Spearman correlation scores of −0.434 for GC3 content and of −0.452 for COUSIN_59_ with p-values of 0.028 and of 0.035). We also find a weak relationship between chromosome size and the number of “Top”, “Middle” and “Bottom” CDSs (Figure 8C). However, these results should be handled with care. Indeed, the Median Absolute Deviation (MAD) scores indicate a broad diversity of GC3 content and COUSIN scores among the studied chromosomes (Supplementary Information 7). Further studies are required to analyse this connection between chromosome size, nucleotide composition and CUPrefs.

**Figure 7:**
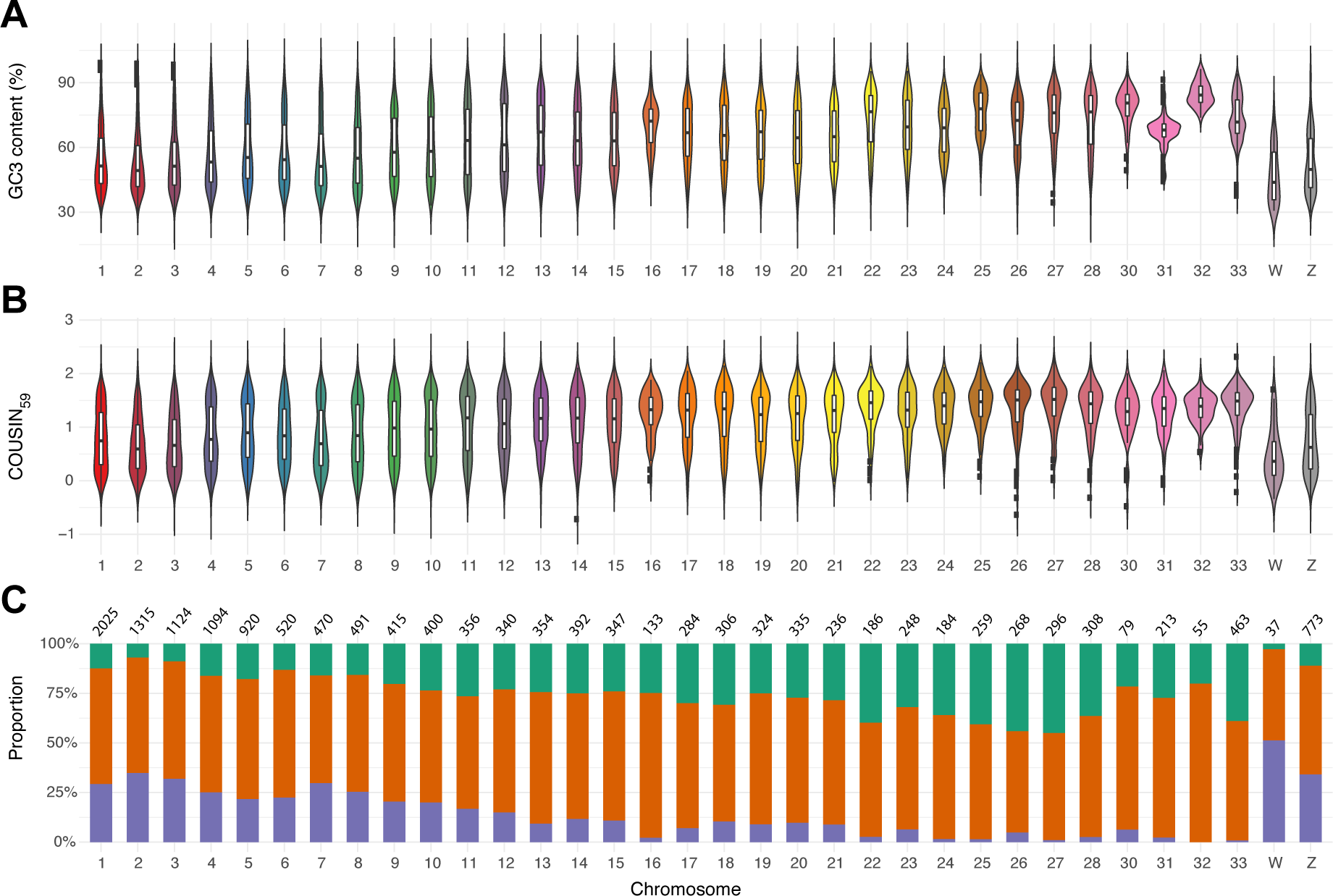
Per chromosome analysis of GC content and COUSIN_59_ scores in CDSs of *G. gallus*. Violin plots of A) GC3 content and B) COUSIN_59_ score along chromosomes of *G. gallus*. C) Proportions of “Top” (green), “Midldle” (orange) and “Bottom” (purple) CDSs in each *G. gallus* chromosomes. The number of CDSs found in each chromosome is shown above each bar.

**Figure 8:**
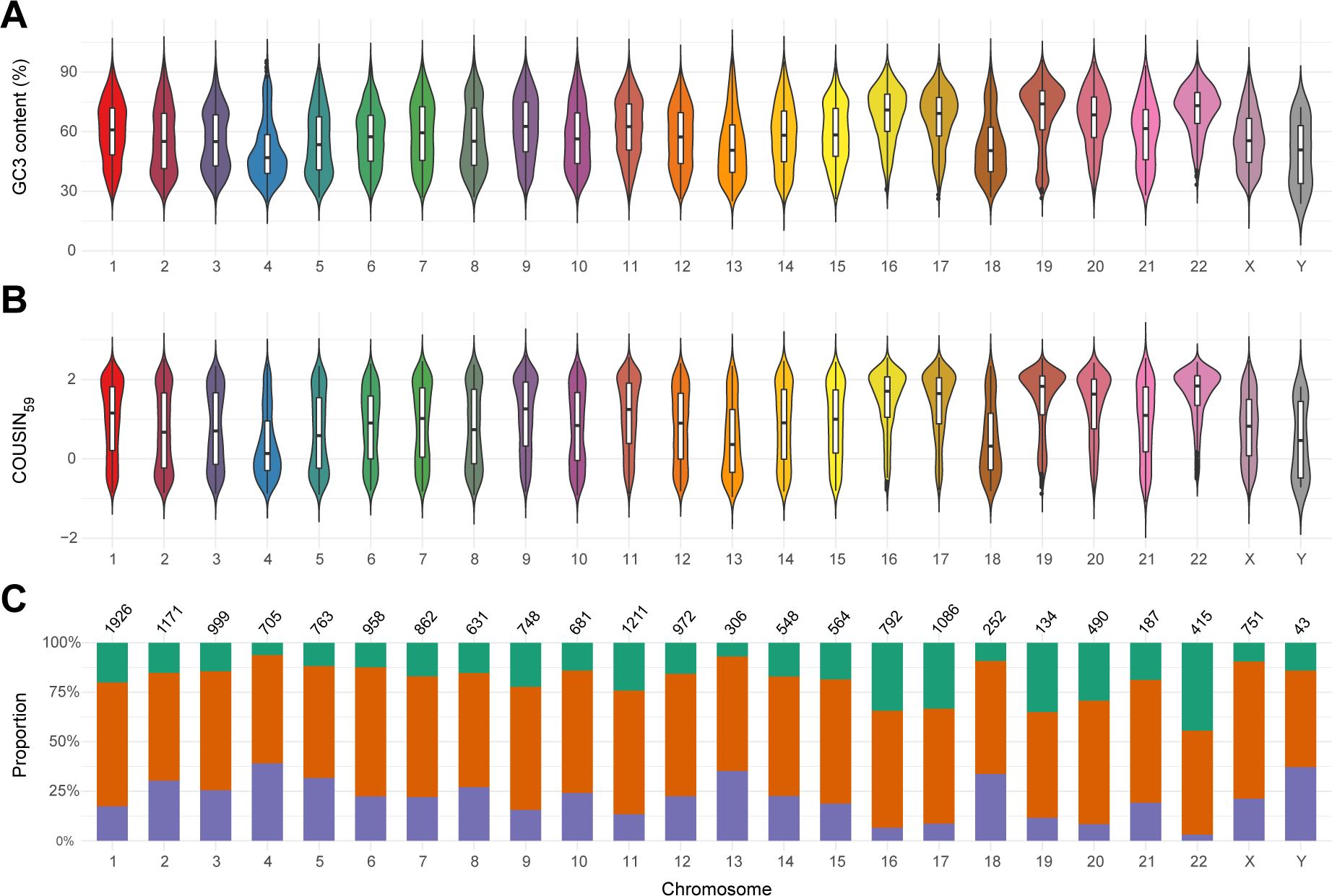
Per chromosome analysis of GC content and COUSIN_59_ scores in CDSs of *H. sapiens*. Violin plots of A) GC3 content and B) COUSIN_59_ score along chromosomes of *H. sapiens*. C) Proportions of “Top” (green), “Midldle” (orange) and “Bottom” (purple) CDSs in each *H. sapiens* chromosomes. The number of CDSs found in each chromosome is shown above each bar.

**Figure 9:**
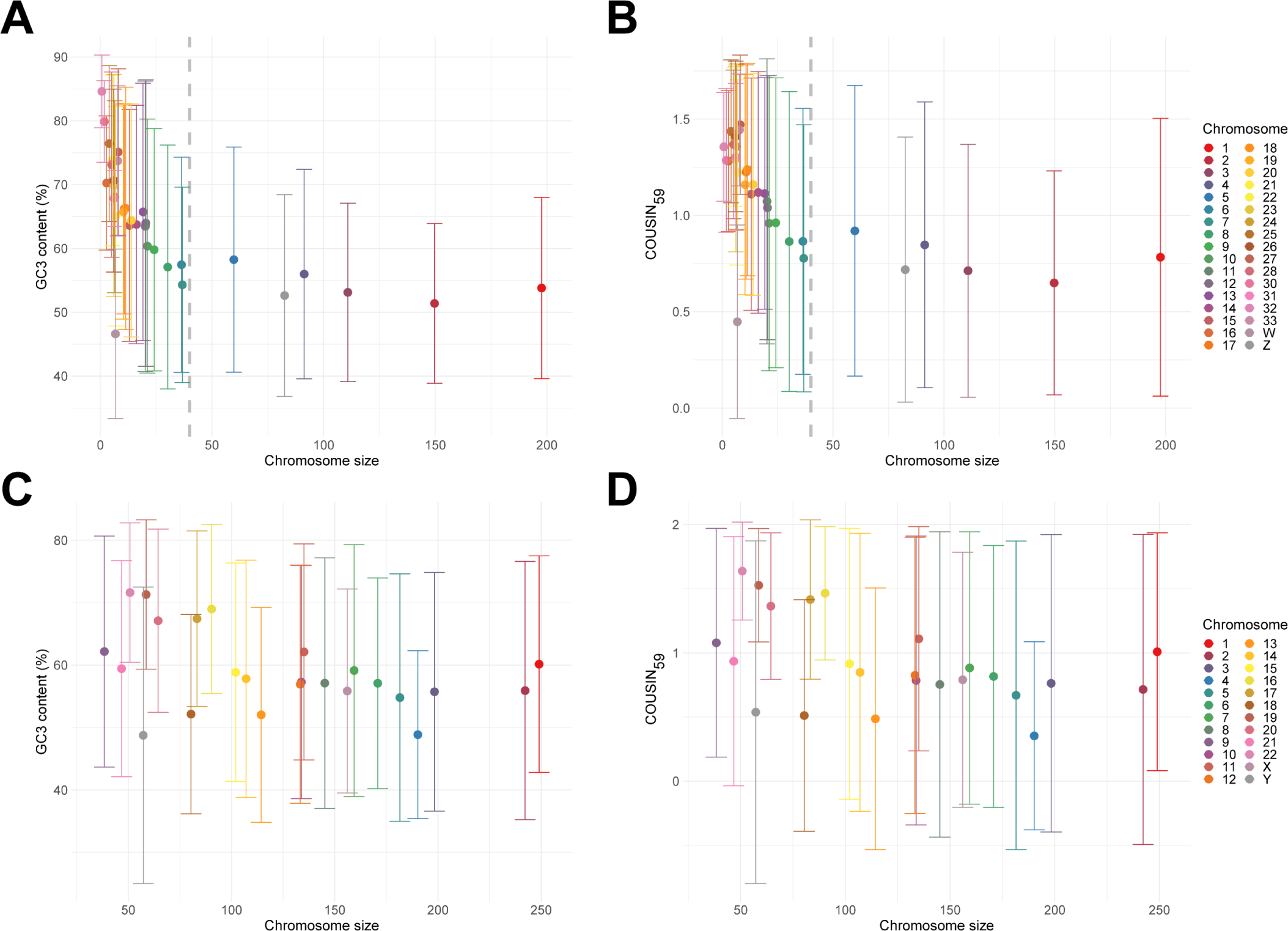
Scatterplots of Huber-M estimator values for GC3 content (A and C) and COUSIN_59_ (B and D) against chromosome size in *G. gallus* (A and B) and *H. sapiens* (C and D). Each dot represents a chromosome with a color indicated in legend. Vertical lines display the MAD values related to the Huber-M estimator ones.

## 5. Discussion

In this study we introduce COUSIN, a new index to measure CUPrefs. This measure has a straightforward quantitative and qualitative meaning, therefore allowing for an easy comparison between the CUPrefs of the query CDS and those of both the reference and a random CUPrefs. We introduce two definitions of the COUSIN index (COUSIN_59_ and COUSIN_18_) depending on whether or not the amino acid composition of the analysed CDS is taken into account when estimating the similarity between its CUPrefs and those of the reference.

We implemented the calculation of the COUSIN index, as well as of a number of additional features and exiting indices to evaluate CUPrefs, in an eponymous bioinformatic software, which is available in a stand-alone and in an online version (`COUSIN`, at http://cousin.ird.fr). This software also estimates confidence intervals of expected COUSIN values given the reference table and the length and composition of the query. Highlighting the limits of these intervals facilitates the evaluation of whether the CUPrefs of the query are significantly different from those expected for a sequence of the same length following the CUPrefs of the reference table.

Finally, we illustrated the novelty and potential of the COUSIN index by applying it to an analysis on eight divergent organisms. During this study, we used the average CUPrefs of the organism as a reference. Importantly, our results show that the use of such average genomic CUPrefs as a sole reference may be more pertinent for certain organisms, such as *P falciparum* or *S. coelicolor*, and less pertinent for other, as exemplified for *H. sapiens* ad *G. gallus*. Using this kind of average reference may or not be relevant when analyzing CUPrefs, as it may allow a first comprehensive understanding of an organism, but may also hide crucial informations on, for example, the CUPrefs of specialized tissues in multicellular organisms. However, the capacity of COUSIN to highlight specificities in organisms while using such reference without preconception shows the strength of this index while analyzing CUPrefs. The analysis of the genomes of two organisms with extreme compositional bias compared to less biased organisms serves to highlight the ease of interpretation of the COUSIN results and the connection between CUPrefs and nucleotide composition. Strikingly, COUSIN unveils a bimodal distributions of CUPrefs on the human and on the chicken genomes hitherto not described. We performed additional analyses that correlate these CUPrefs bimodal distributions with the GC3 distribution in these genomes as well as the specific genomic context of the corresponding CDSs in terms of chromosomal location.

Overall, the novel COUSIN index and COUSIN software can serve as an intuitive and powerful software to analyse CUPrefs. Our results on the human genome will undoubtedly foster new research on the mutation-selection dynamics that pattern CUPrefs.

## Supporting information

All supplemental data

## 6. ACKNOWLEDGEMENTS

Jérôme Bourret is funded by a PhD fellowship from the *French Ministry of Education and Research*. This work was supported by the European Union’s Horizon 2020 research and innovation programme under the grant agreement CODOVIREVOL (ERC-2014-CoG-647916) to IGB. The authors acknowledge the CNRS and the IRD for additional support. The computational results presented have been achieved in part using the IRD Bioinformatic Cluster *itrop*, which also hosts the COUSIN online server (http://cousin.ird.fr). We thank Frederic Delsuc (ISEM, Université de Montpellier) for driving our attention onto the compositional peculiarities of the *G. gallus* genome, that have served to illustrate our analysis of CUPrefs in organisms with micro and macrochromosomes.

