## Supplementary material for "COUSIN (COdon Usage Similarity INdex): A normalized measure of Codon Usage Preferences": All supplemental data

### 1 CAI calculation

the first step in the CAI index as described by Sharp et Li Sharp and Li (1987) consists in defining for each synonymous codon of an amino acid a relative adaptiveness score ($w_{c,a}$). In the reference dataset, this value is obtained by calculating the ratio between the frequency of a codon ($f_{c,a}^{\text{ref}}$) and the frequency of its most represented synonymous ($f_{\text{cmax,a}}^{\text{ref}}$):

$$\mathrm{w_{c,a}} = \frac{f_{c,a}^{\text{ref}}}{f_{\text{cmax,a}}^{\text{ref}}}$$

The CAI score is obtained by calculating the geoindexal mean of the relative adaptivenesses multiplied by the occurrences of the related codons found in the query sequence (Sharp and Li, 1987). Here, we name this CAI score $\text{CAI}_{59}$, since all 59 synonymous codons have an impact on the CAI score regardless of their relation with their amino acid:

$$\text{CAI}_{59} = \left( \prod_{a \in \mathcal{A}} \prod_{c \in k_a} \text{Occ}_{c,a}^{\text{que}} \times w_{c,a} \right)^{\frac{1}{L}}$$

where $\text{Occ}_{c,a}^{\text{que}}$ is the number of occurrences of codon $c$ in the query and $L$ is the length of the query (total number of amino acids).

### 2 Average % of G+C content in studied organisms

The average % of G+C content in the 8 organisms analyzed are given in Table 1.

Table 1: Average % of GC3 content of complete CDSs of *Escherichia coli*, *Streptomyces coelicolor*, *Saccharomyces cerevisiae*, *plasmodium falciparum*, *Homo sapiens*, *Mus musculus* and *Arabidopsis thaliana*. This table describes the average GC3 content found among all CDSs, but also in CDSs with the top 20%, the 60% in the middle and the bottom 20% $\text{COUSIN}_{59}$ score.

| | ***E. coli*** | ***S. coelicolor*** | ***S. cerevisiae*** | ***P. falciparum*** | ***H. sapiens*** | ***G. gallus*** | ***M. musculus*** | ***A. thaliana*** |
|---|---|---|---|---|---|---|---|---|
| **GC3 content (all dataset)** | 54.906 | 92.373 | 39.211 | 17.797 | 59.977 | 60.635 | 58.641 | 42.717 |
| **GC3 content (top 20%)** | 59.976 | 96.621 | 33.976 | 13.910 | 78.971 | 77.917 | 71.825 | 37.806 |
| **GC3 content (middle 60%)** | 56.128 | 93.517 | 37.976 | 17.325 | 61.056 | 61.236 | 59.537 | 42.114 |
| **GC3 content (bottom 20%)** | 46.169 | 84.693 | 48.146 | 23.097 | 37.739 | 41.549 | 42.771 | 49.436 |

### 3 Global analysis of CUPrefs scores against GC content in organisms

In addition to the first analysis putting on sight the dispersion of COUSIN and CAI scores among organisms CDSs, we also draw up the scores of these same CDSs with their GC3 content to bring light on the relation between these two variables. Figures 1, 2 ($\text{COUSIN}_{59}$), 3 and 4 ($\text{CAI}_{59}$) show individual scatterplots between one of the two index score and the GC3 content along with Pearson correlation tests.

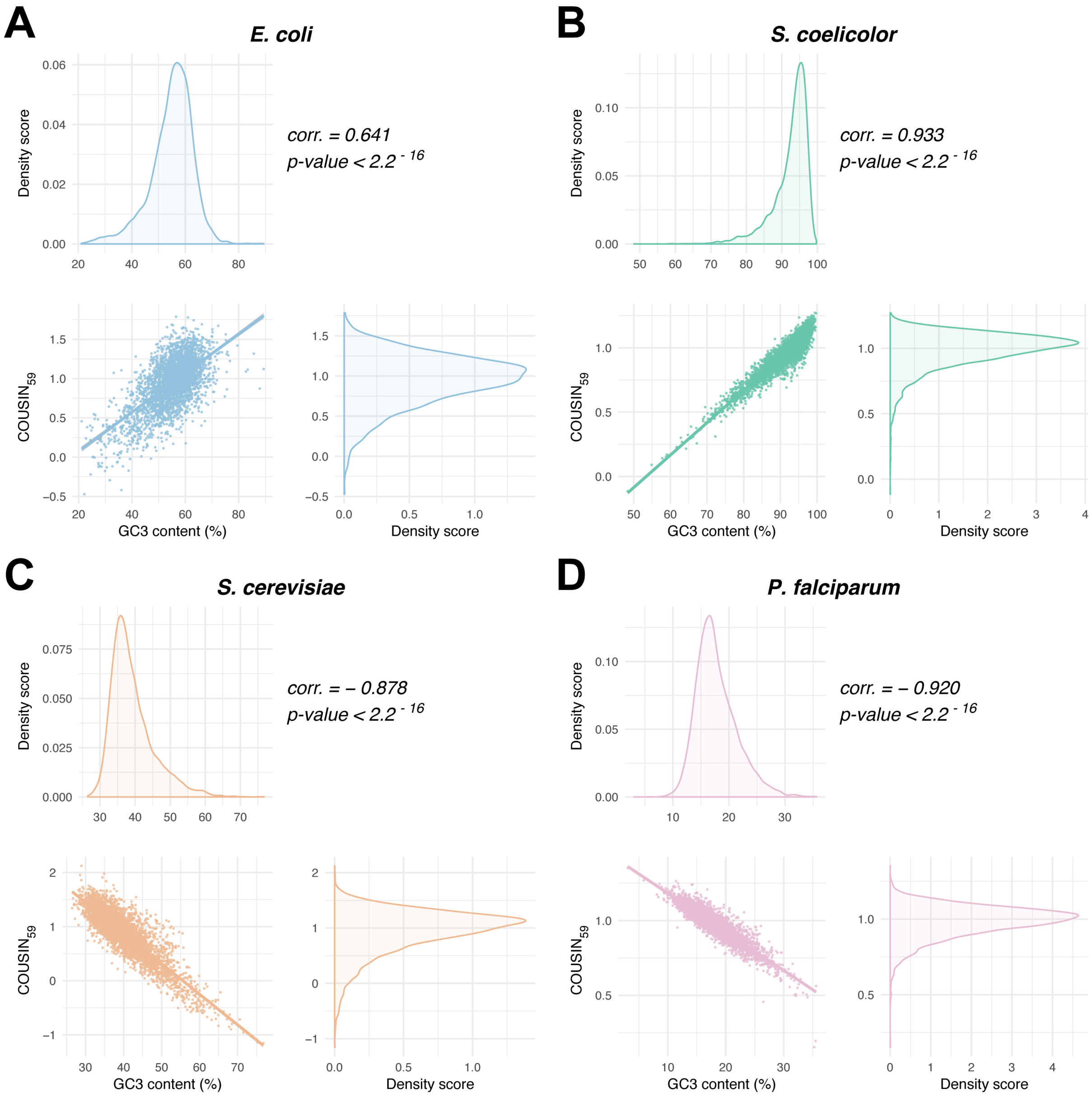


Figure 1: Scatterplot of $\text{COUSIN}_{59}$ (x-axis) and GC3 content (y-axis) scores for CDSs belonging to *E. coli* (A), *S. coelicolor* (B), *S. cerevisiae* (C) and *P. falciparum* (D). Each scatterplot is accompanied by two density curves: $\text{COUSIN}_{59}$ (right of the scatterplot) and GC3 content (top of the scatterplot). On the top-right of the scatterplots, statistics of Pearson correlation tests between $\text{COUSIN}_{59}$ scores and GC3 content is given.

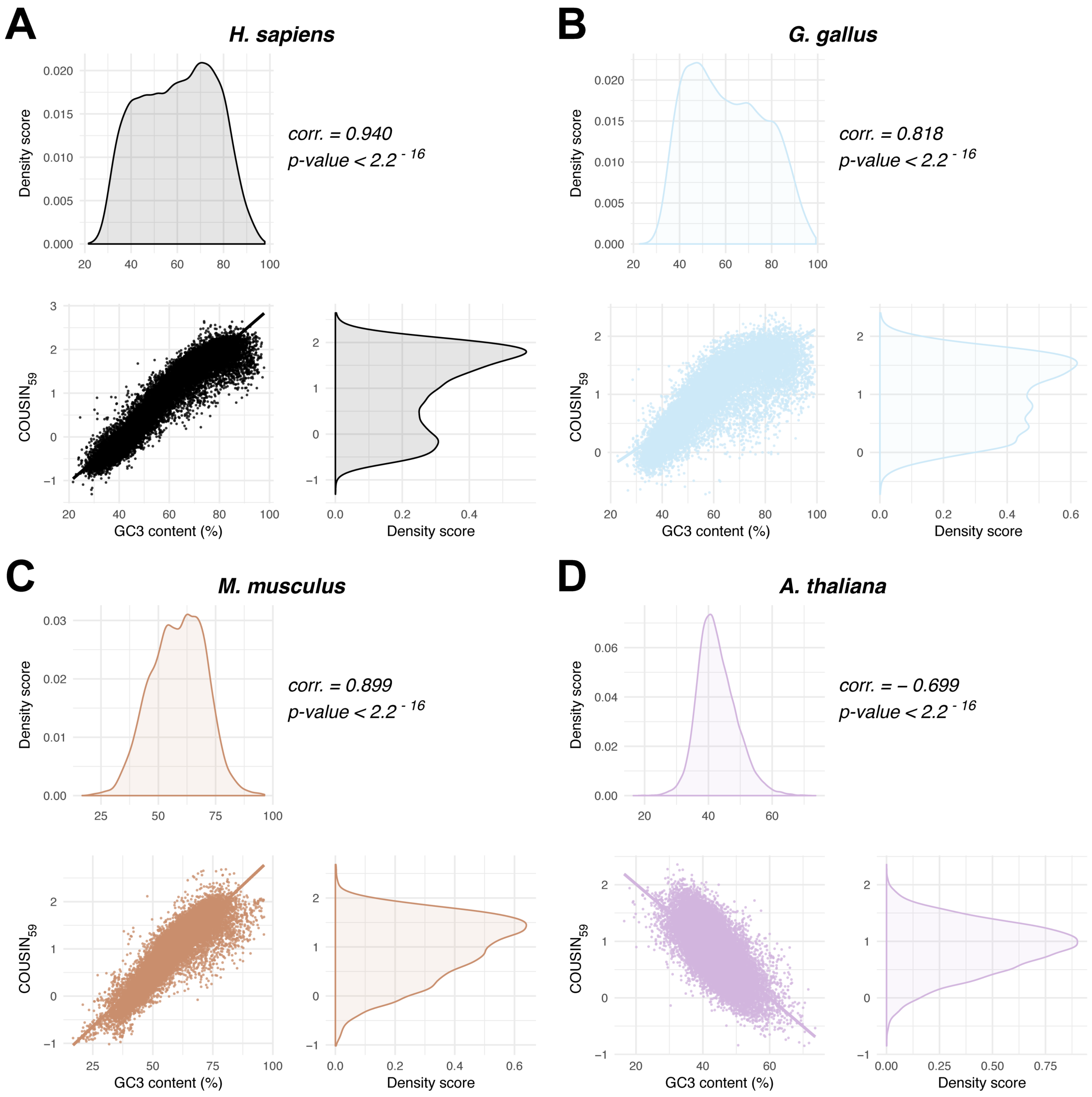


Figure 2: Scatterplot of $COUSIN_{59}$ (x-axis) and GC3 content (y-axis) scores for CDSs belonging to *H. sapiens* (A), *G. gallus* (B), *Mus musculus* (C) and *A. thaliana* (D). Each scatterplot is accompanied by two density curves: $COUSIN_{59}$ (right of the scatterplot) and GC3 content (top of the scatterplot). On the top-right of the scatterplots, statistics of Pearson correlation tests between $COUSIN_{59}$ scores and GC3 content is given.

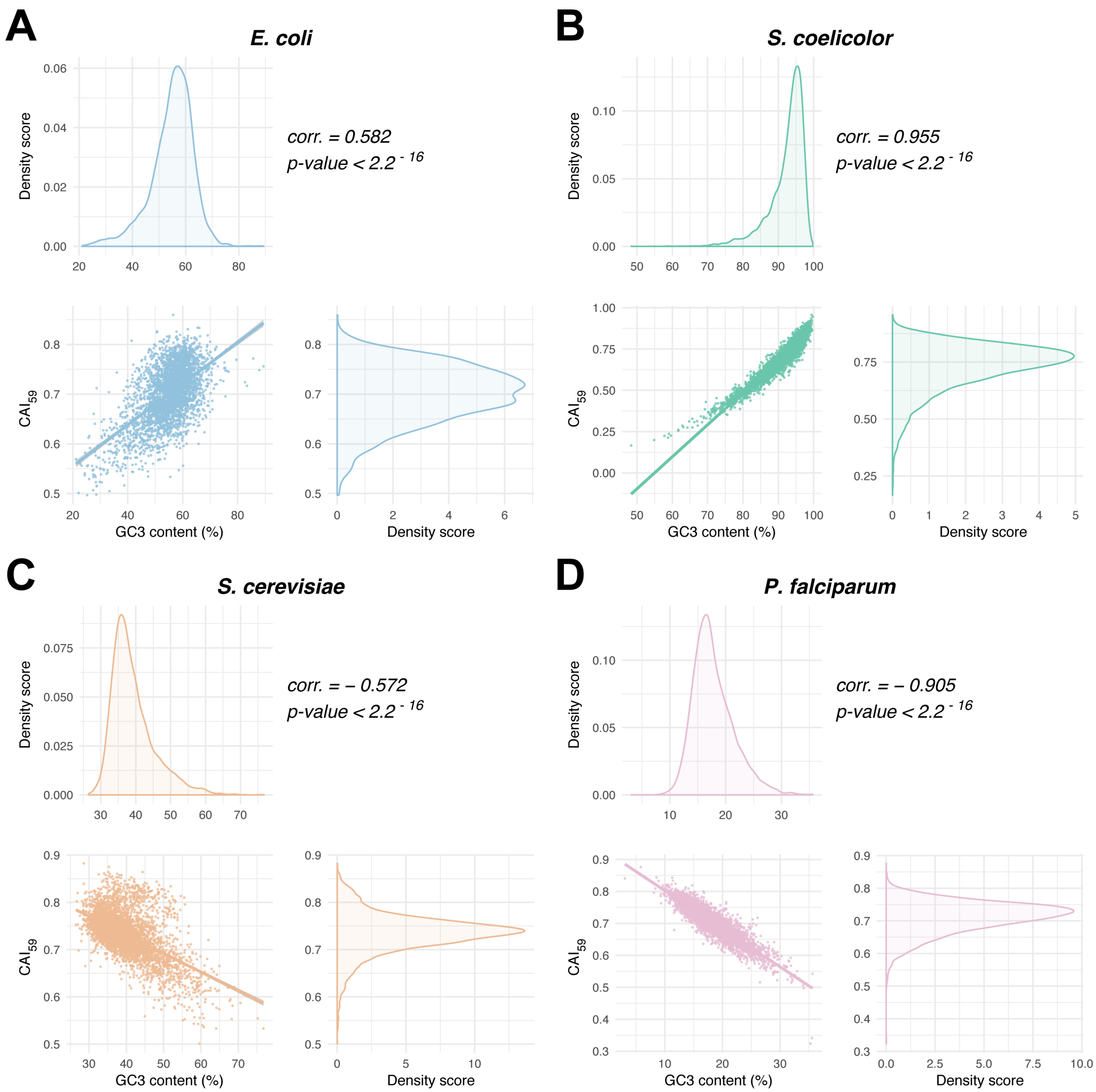


Figure 3: Scatterplot of $CAI_{59}$ (x-axis) and GC3 content (y-axis) scores for CDSs belonging to *E. coli* (A), *S. coelicolor* (B), *S. cerevisiae* (C) and *P. falciparum* (D). Each scatterplot is accompanied by two density curves: $CAI_{59}$ (right of the scatterplot) and GC3 content (top of the scatterplot). On the top-right of the scatterplots, statistics of Pearson correlation tests between $CAI_{59}$ scores and GC3 content is given.

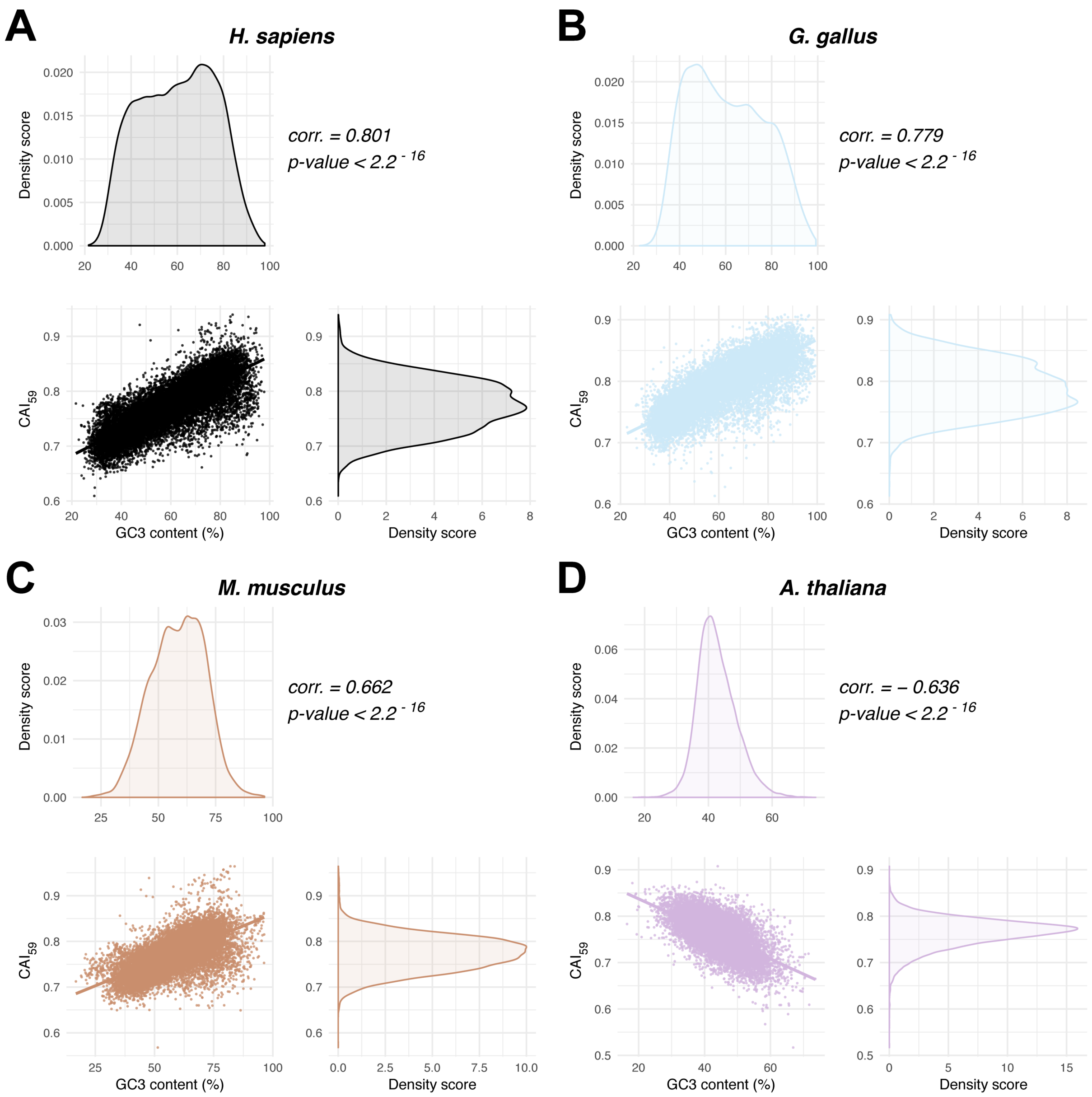


Figure 4: Scatterplot of $CAI_{59}$ (x-axis) and GC3 content (y-axis) scores for CDSs belonging to *H. sapiens* (A), *G. gallus* (B), *Mus musculus* (C) and *A. thaliana* (D). Each scatterplot is accompanied by two density curves: $CAI_{59}$ (right of the scatterplot) and GC3 content (top of the scatterplot). On the top-right of the scatterplots, statistics of Pearson correlation tests between $CAI_{59}$ scores and GC3 content of CDSs is given.

#### 4 Statistics on studied organisms

The mean values, Huber-M estimator values and MAD scores related to the CAI and COUSIN analysis on the studied organisms are given in Table 2.

Table 2: Mean value, Huber-M estimator value and MAD scores of $COUSIN_{18}$, $COUSIN_{59}$, $CAI_{18}$ and $CAI_{59}$ scores on the studied organisms.

| | ***E. coli*** | ***S. coelicolor*** | ***S. cerevisiae*** | ***P. falciparum*** | ***H. sapiens*** | ***G. gallus*** | ***M. musculus*** | ***A. thaliana*** |
|---|---|---|---|---|---|---|---|---|
| **Mean (COUSIN**$_{18}$**)** | 0.928 | 0.980 | 0.909 | 0.975 | 0.975 | 0.990 | 0.982 | 0.849 |
| **Huber-M estimator (COUSIN**$_{18}$**)** | 0.936 | 0.995 | 0.949 | 0.984 | 0.975 | 0.991 | 0.991 | 0.859 |
| **MAD (COUSIN**$_{18}$**)(+/-)** | 0.331 | 0.102 | 0.334 | 0.095 | 1.229 | 0.851 | 0.808 | 0.498 |
| **Mean (COUSIN**$_{59}$**)** | 0.942 | 0.981 | 0.925 | 0.982 | 0.949 | 0.968 | 0.967 | 0.868 |
| **Huber-M estimator (COUSIN**$_{59}$**)** | 0.959 | 0.996 | 0.967 | 0.989 | 0.951 | 0.969 | 0.982 | 0.877 |
| **MAD (COUSIN**$_{59}$**)(+/-)** | 0.285 | 0.111 | 0.315 | 0.088 | 1.026 | 0.740 | 0.686 | 0.451 |
| **Mean (CAI**$_{18}$**)** | 0.737 | 0.765 | 0.741 | 0.707 | 0.809 | 0.820 | 0.808 | 0.785 |
| **Huber-M estimator (CAI**$_{18}$**)** | 0.740 | 0.777 | 0.742 | 0.710 | 0.809 | 0.230 | 0.808 | 0.787 |
| **MAD (CAI**$_{18}$**)(+/-)** | 0.046 | 0.088 | 0.031 | 0.046 | 0.053 | 0.048 | 0.035 | 0.023 |
| **Mean (CAI**$_{59}$**)** | 0.700 | 0.726 | 0.734 | 0.710 | 0.773 | 0.790 | 0.773 | 0.764 |
| **Huber-M estimator (CAI**$_{59}$**)** | 0.702 | 0.736 | 0.736 | 0.713 | 0.773 | 0.790 | 0.773 | 0.765 |
| **MAD (CAI**$_{59}$**)(+/-)** | 0.058 | 0.087 | 0.032 | 0.043 | 0.051 | 0.047 | 0.039 | 0.027 |

### 5 Global analysis of correlation between COUSIN and CAI indexes

Scatter-plots along with Pearson correlation tests between COUSIN and CAI indexes are given in Figures 5 (*E. coli*), 6 (*S. coelicolor*), 7 (*S. cerevisiae*), 8 (*P. falciparum*), 9 (*H. sapiens*), 10 (*G. gallus*), 11 (*M. musculus*), 12 (*A. thaliana*).

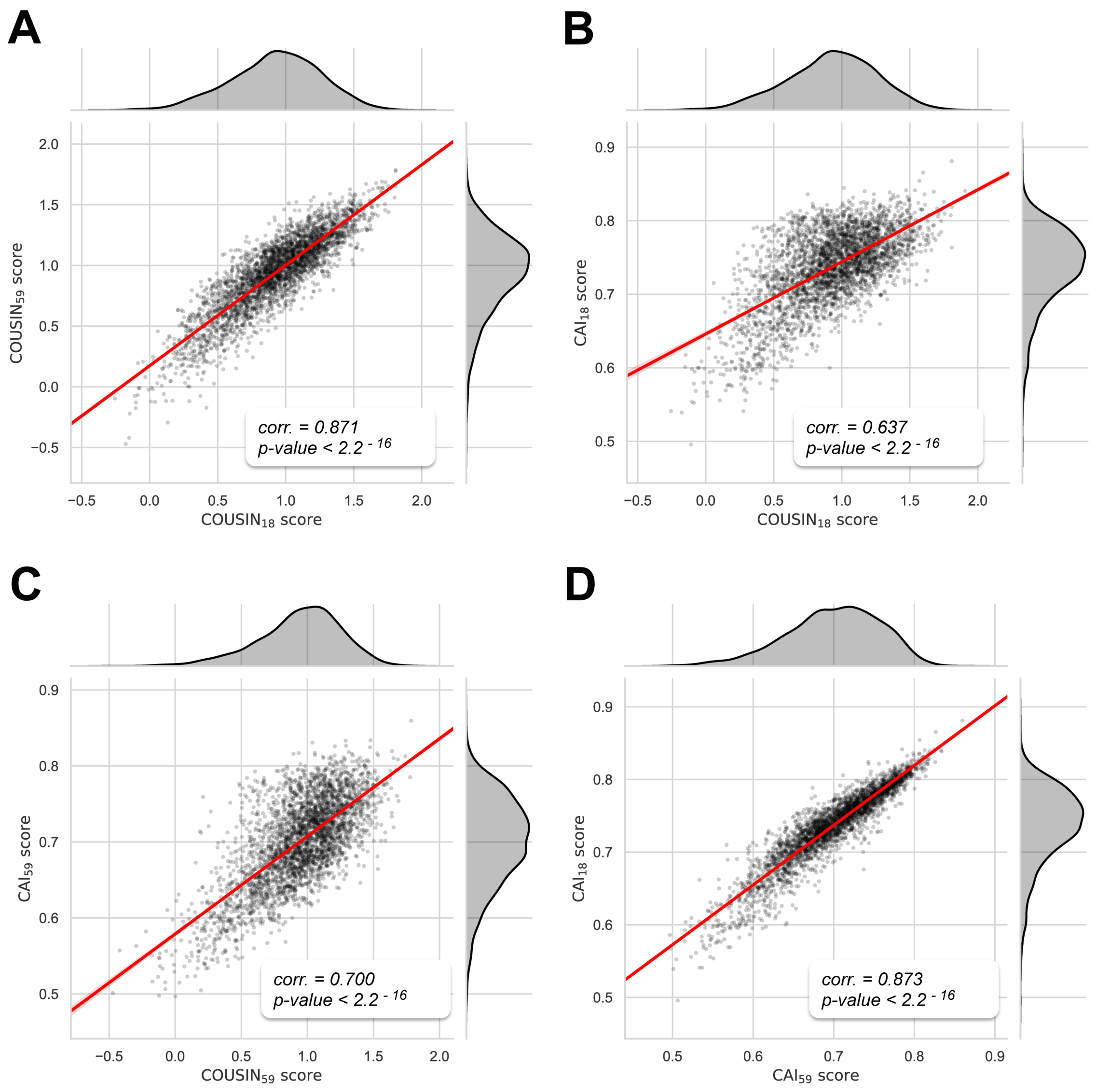


Figure 5: Dot-plots of *E. coli* CDSs scores between COUSIN metrics declinations, CAI metrics declinations and between both metrics. In addition to the dot-plot, a blue regression line is given with its 95% confidence interval. For each plot, the x-axis and y-axis represent scores obtained for one metric. Results of Pearson's correlation test are indicated on the top-right of plots. Histograms and density plots are given at the opposite of x-axis and y-axis legends. These additional plots indicate the distribution of scores with the related index.

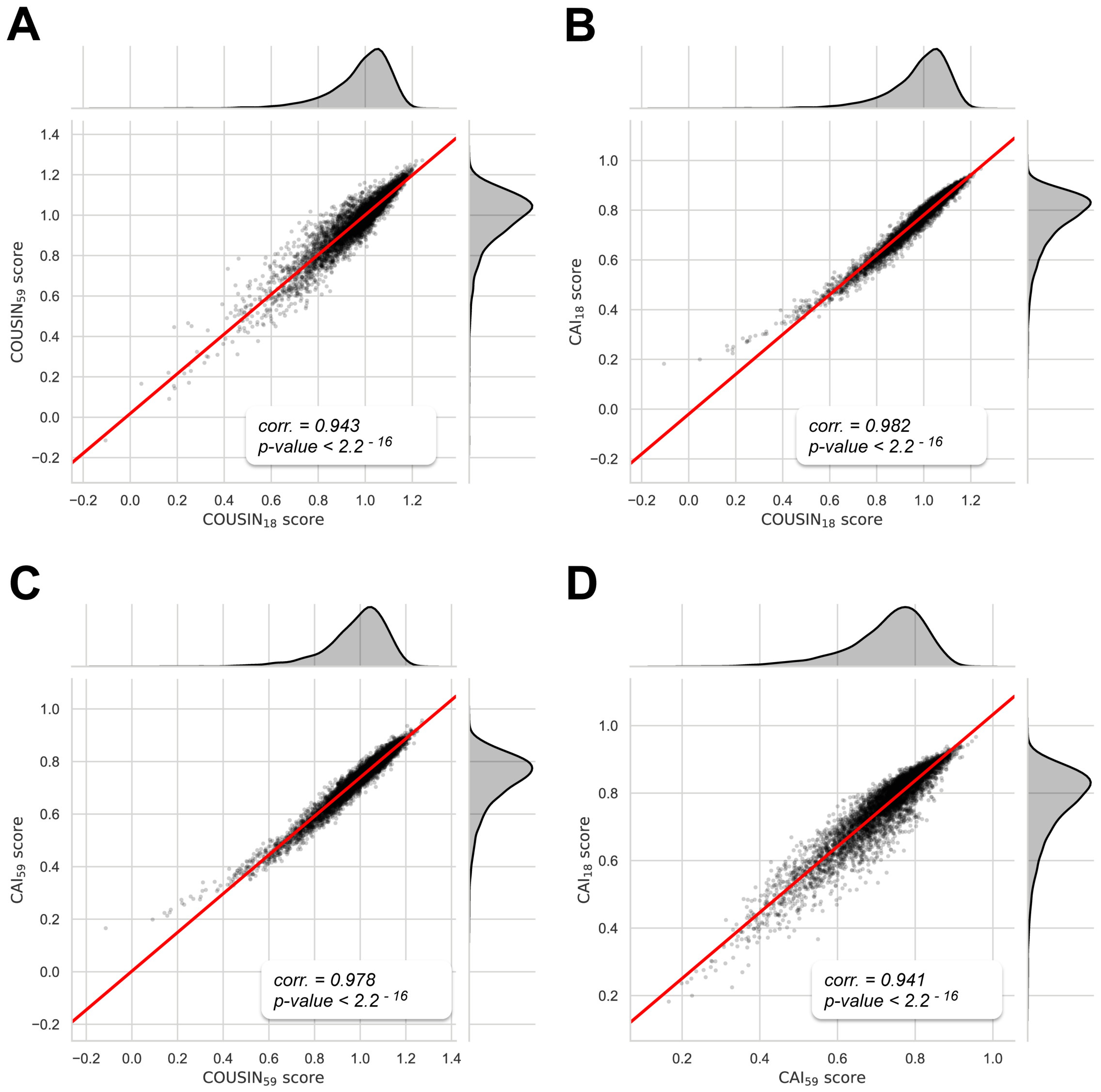


Figure 6: Dot-plots of *S. coelicolor* CDSs scores between $\text{COUSIN}_{18}$ and $\text{COUSIN}_{59}$ (A),$\text{COUSIN}_{18}$ and $\text{CAI}_{18}$ (B), $\text{COUSIN}_{59}$ and $\text{CAI}_{59}$ (C) and between $\text{CAI}_{18}$ and $\text{CAI}_{59}$ indexes (D). In addition to the dot-plot, a blue regression line is given with its 95% confidence interval. For each plot, the x-axis and y-axis represent scores obtained for one metric. Results of Pearson's correlation test are indicated on the top-right of plots. Histograms and density plots are given at the opposite of x-axis and y-axis legends. These additional plots indicate the distribution of scores with the related index.

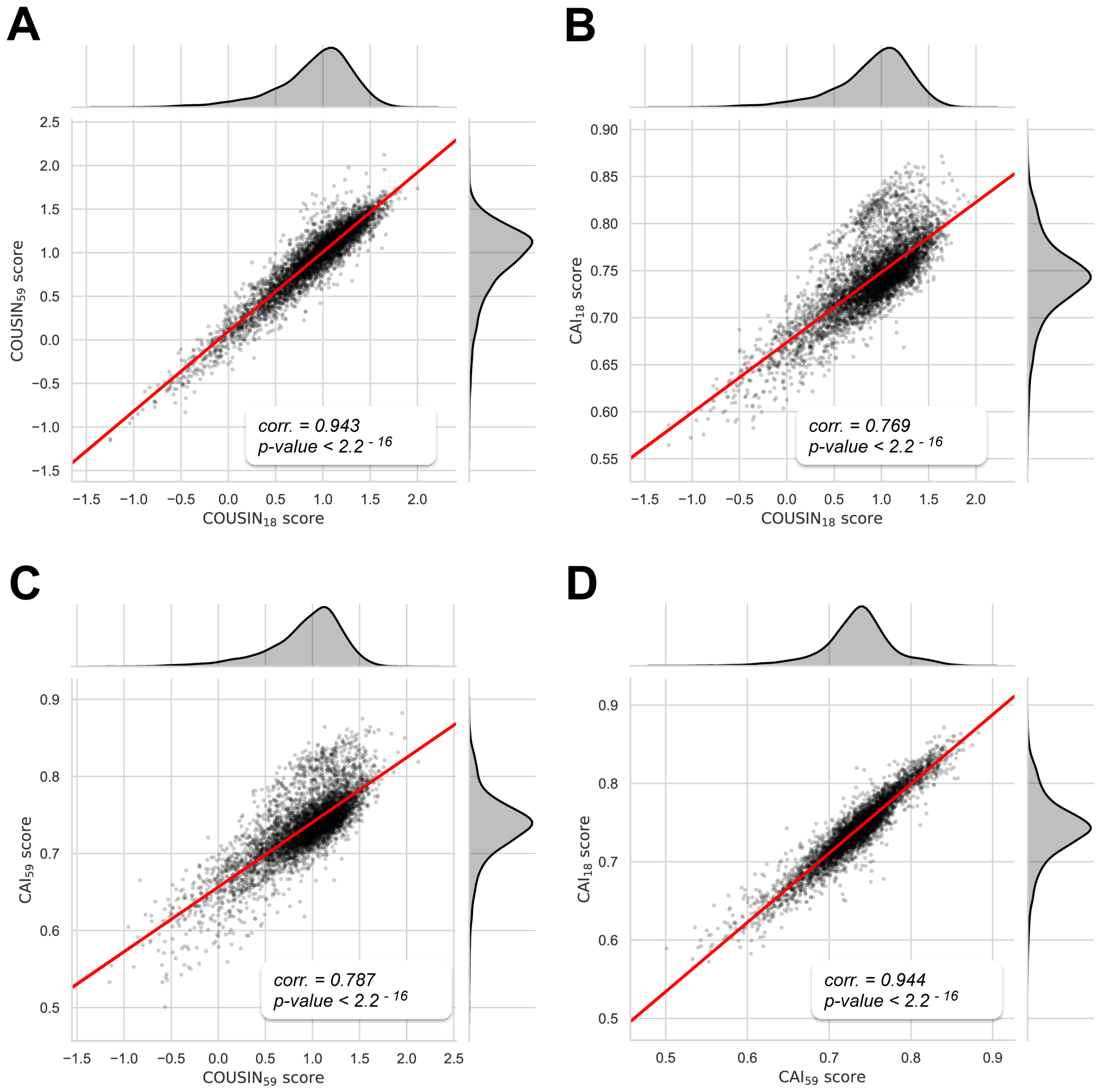


Figure 7: Dot-plots of *S. cerevisiae* CDSs scores between $\text{COUSIN}_{18}$ and $\text{COUSIN}_{59}$ (A),$\text{COUSIN}_{18}$ and $\text{CAI}_{18}$ (B), $\text{COUSIN}_{59}$ and $\text{CAI}_{59}$ (C) and between $\text{CAI}_{18}$ and $\text{CAI}_{59}$ indexes (D). In addition to the dot-plot, a blue regression line is given with its 95% confidence interval. For each plot, the x-axis and y-axis represent scores obtained for one metric. Results of Pearson's correlation test are indicated on the top-right of plots. Histograms and density plots are given at the opposite of x-axis and y-axis legends. These additional plots indicate the distribution of scores with the related index.

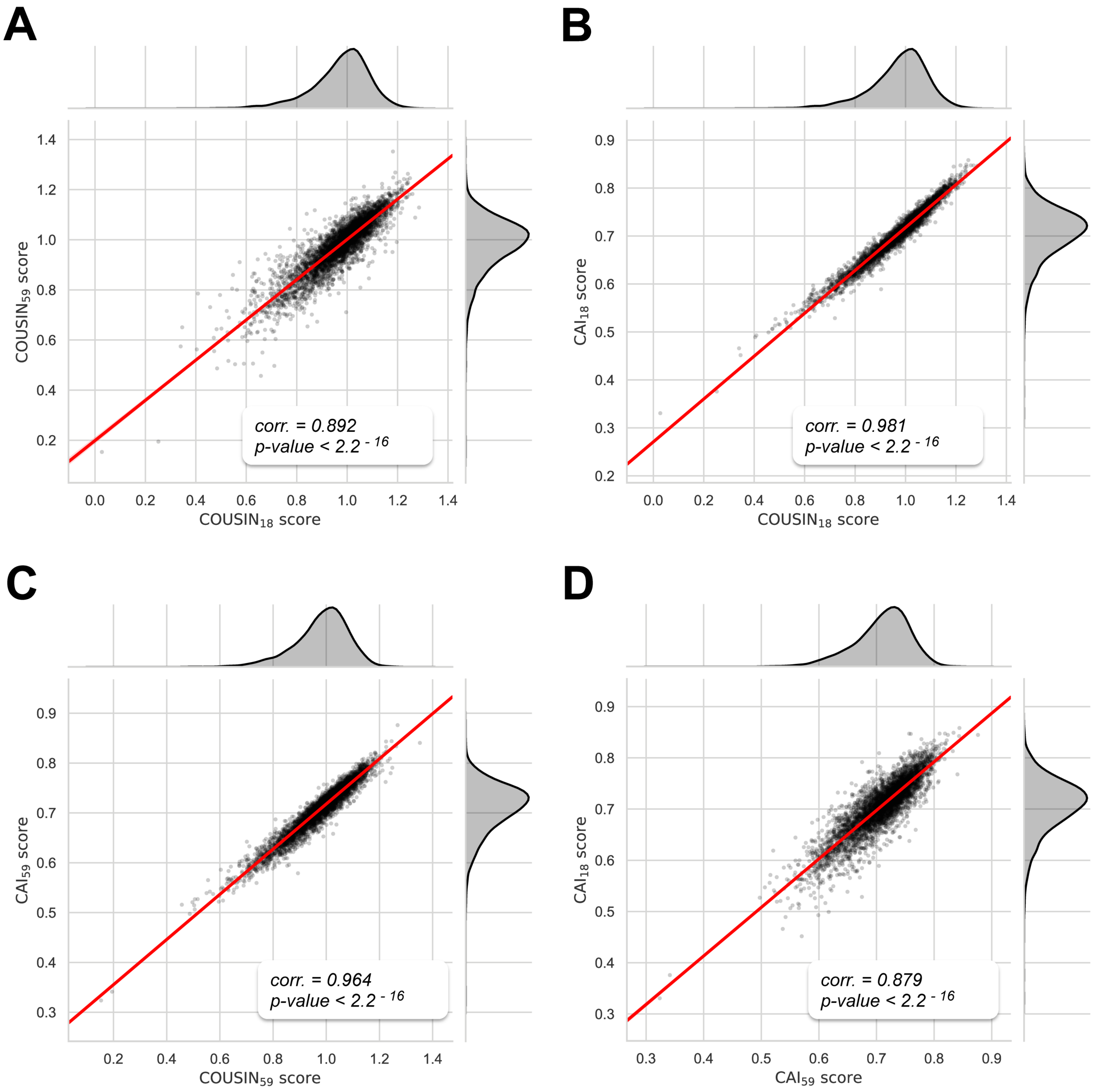


Figure 8: Dot-plots of *P. falciparum* CDSs scores between $\mathrm{COUSIN}_{18}$ and $\mathrm{COUSIN}_{59}$ (A),$\mathrm{COUSIN}_{18}$ and $\mathrm{CAI}_{18}$ (B), $\mathrm{COUSIN}_{59}$ and $\mathrm{CAI}_{59}$ (C) and between $\mathrm{CAI}_{18}$ and $\mathrm{CAI}_{59}$ indexes (D). In addition to the dot-plot, a blue regression line is given with its 95% confidence interval. For each plot, the x-axis and y-axis represent scores obtained for one metric. Results of Pearson's correlation test are indicated on the top-right of plots. Histograms and density plots are given at the opposite of x-axis and y-axis legends. These additional plots indicate the distribution of scores with the related index.

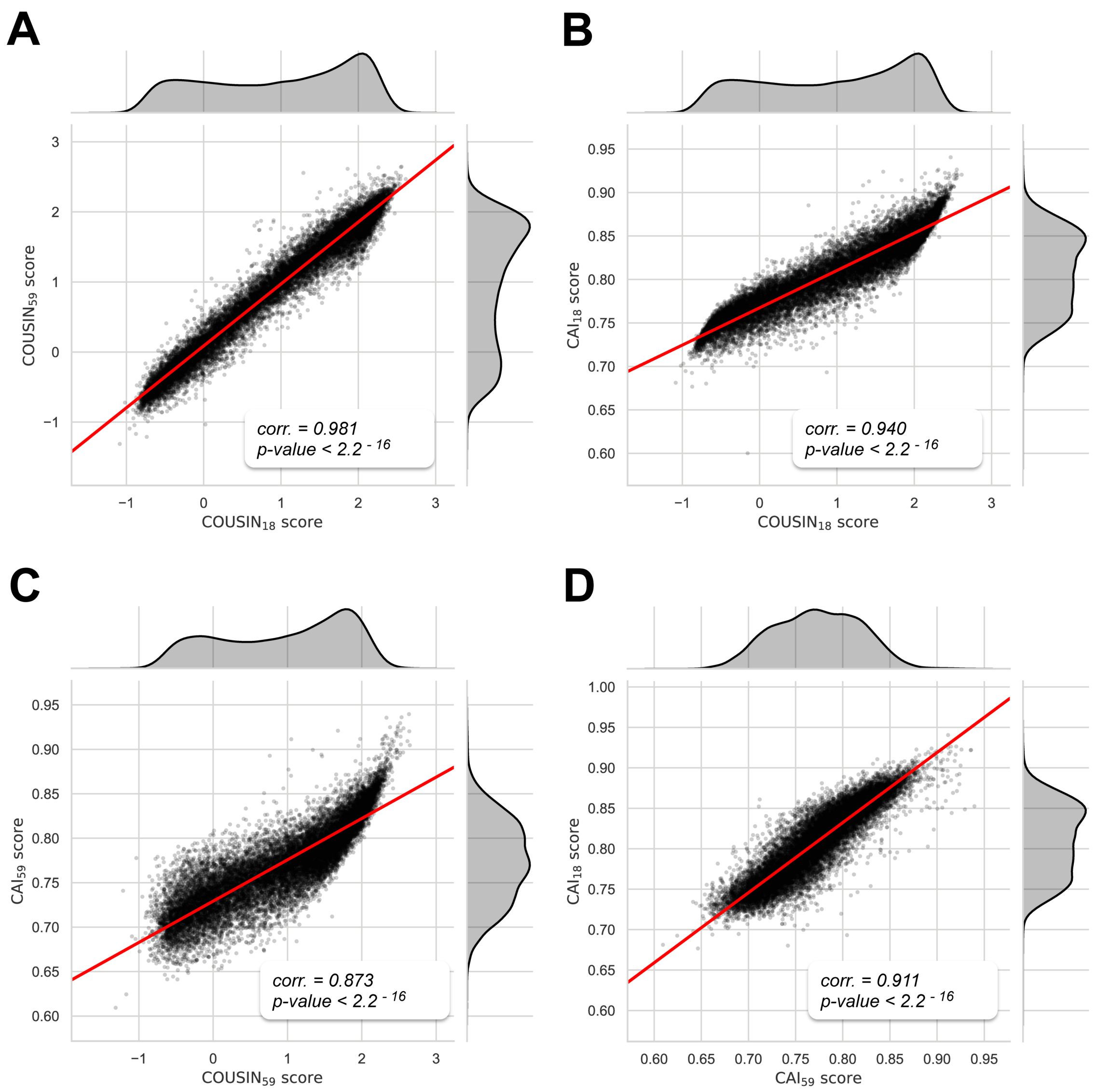


Figure 9: Dot-plots of *H. sapiens* CDSs scores between $COUSIN_{18}$ and $COUSIN_{59}$ (A),$COUSIN_{18}$ and $CAI_{18}$ (B), $COUSIN_{59}$ and $CAI_{59}$ (C) and between $CAI_{18}$ and $CAI_{59}$ indexes (D). In addition to the dot-plot, a blue regression line is given with its 95% confidence interval. For each plot, the x-axis and y-axis represent scores obtained for one metric. Results of Pearson's correlation test are indicated on the top-right of plots. Histograms and density plots are given at the opposite of x-axis and y-axis legends. These additional plots indicate the distribution of scores with the related index.

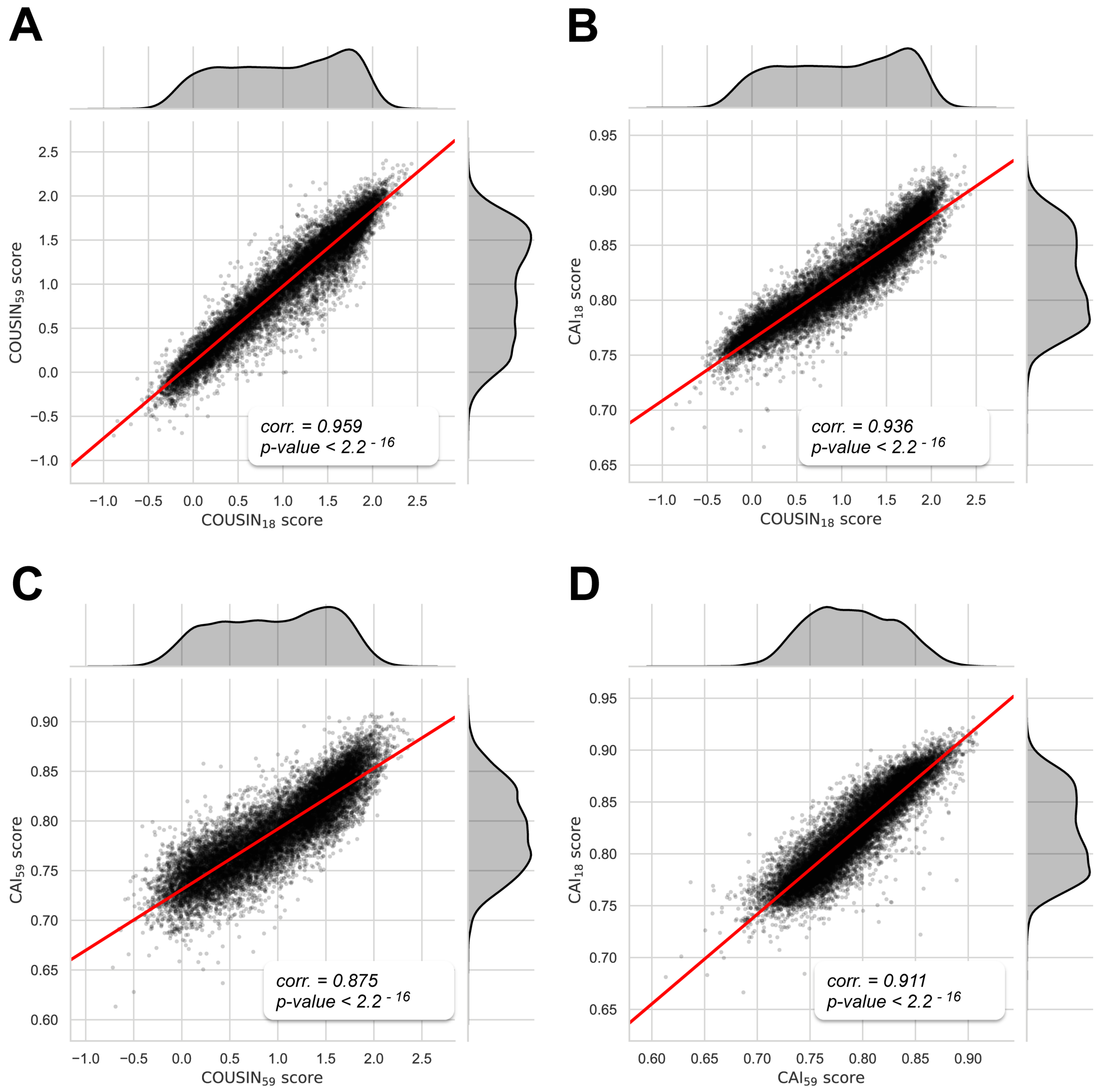


Figure 10: Dot-plots of *G. gallus* CDSs scores between COUSIN$_{18}$ and COUSIN$_{59}$ (A),COUSIN$_{18}$ and CAI$_{18}$ (B), COUSIN$_{59}$ and CAI$_{59}$ (C) and between CAI$_{18}$ and CAI$_{59}$ indexes (D). In addition to the dot-plot, a blue regression line is given with its 95% confidence interval. For each plot, the x-axis and y-axis represent scores obtained for one metric. Results of Pearson's correlation test are indicated on the top-right of plots. Histograms and density plots are given at the opposite of x-axis and y-axis legends. These additional plots indicate the distribution of scores with the related index.

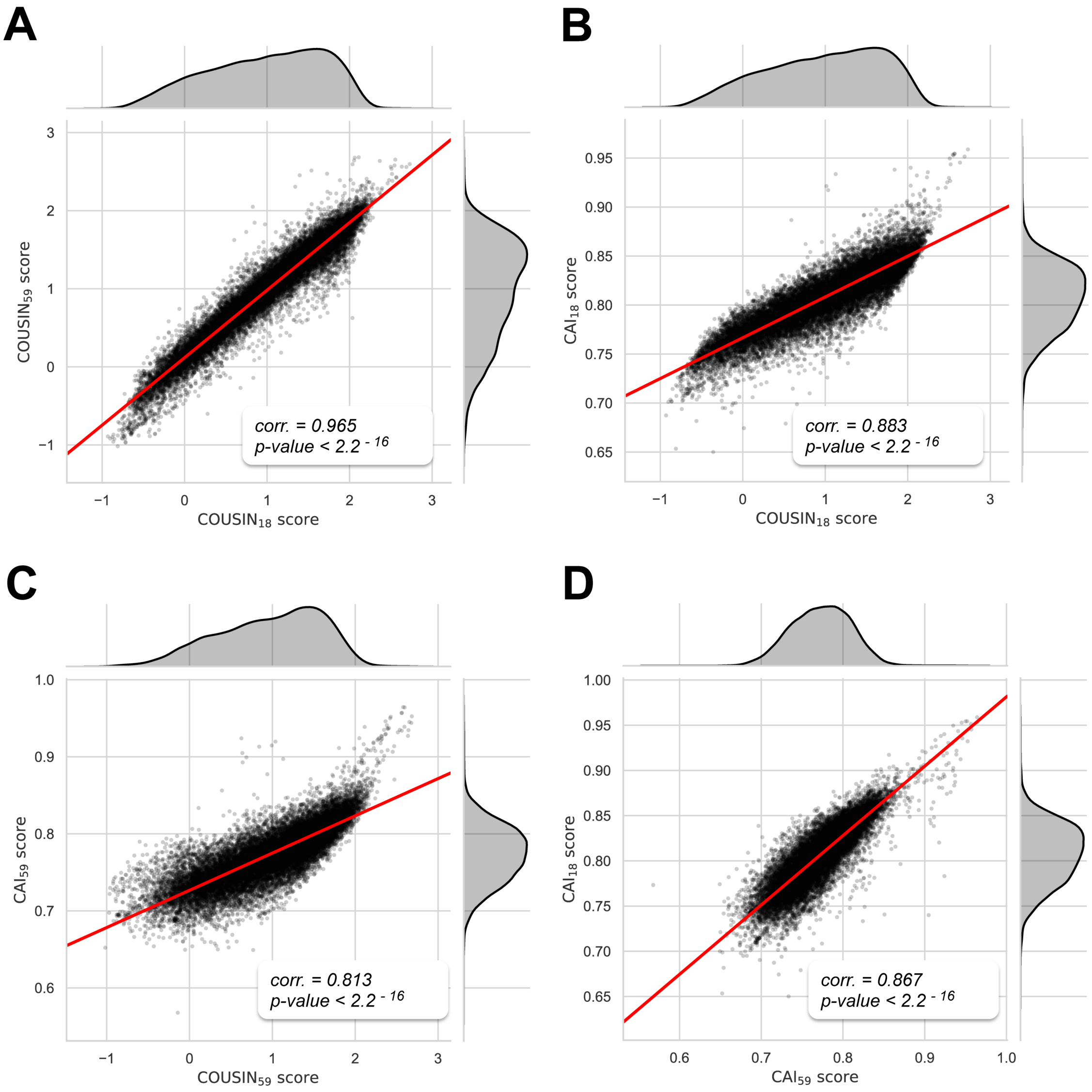


Figure 11: Dot-plots of *M. musculus* CDSs scores between $COUSIN_{18}$ and $COUSIN_{59}$ (A),$COUSIN_{18}$ and $CAI_{18}$ (B), $COUSIN_{59}$ and $CAI_{59}$ (C) and between $CAI_{18}$ and $CAI_{59}$ indexes (D). In addition to the dot-plot, a blue regression line is given with its 95% confidence interval. For each plot, the x-axis and y-axis represent scores obtained for one metric. Results of Pearson's correlation test are indicated on the top-right of plots. Histograms and density plots are given at the opposite of x-axis and y-axis legends. These additional plots indicate the distribution of scores with the related index.

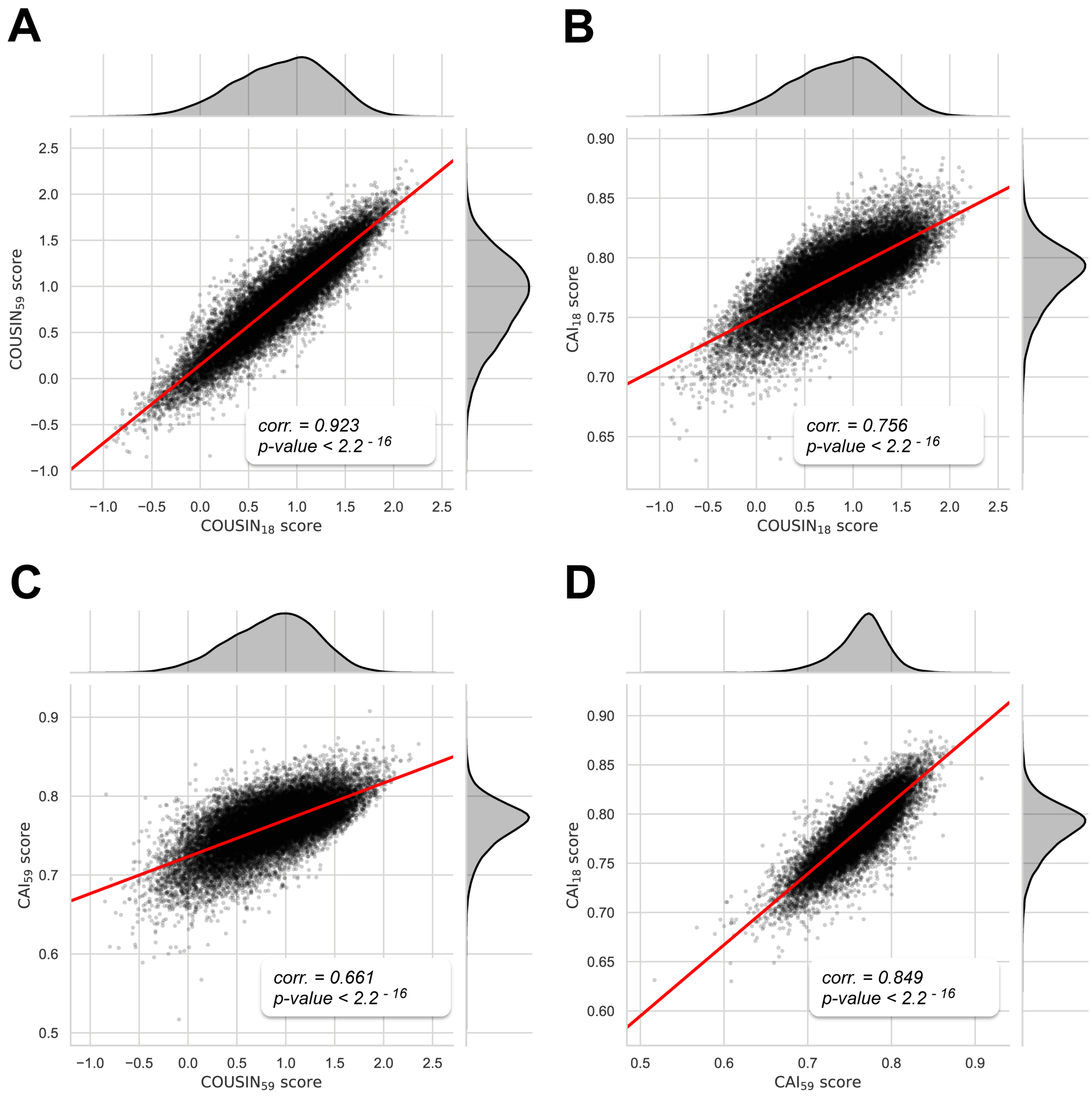


Figure 12: Dot-plots of *A. thaliana* CDSs scores between $\text{COUSIN}_{18}$ and $\text{COUSIN}_{59}$ (A),$\text{COUSIN}_{18}$ and $\text{CAI}_{18}$ (B), $\text{COUSIN}_{59}$ and $\text{CAI}_{59}$ (C) and between $\text{CAI}_{18}$ and $\text{CAI}_{59}$ indexes (D). In addition to the dot-plot, a blue regression line is given with its 95% confidence interval. For each plot, the x-axis and y-axis represent scores obtained for one metric. Results of Pearson's correlation test are indicated on the top-right of plots. Histograms and density plots are given at the opposite of x-axis and y-axis legends. These additional plots indicate the distribution of scores with the related index.

#### 6 CUPrefs anaysis on *H. sapiens* chromosomes

Intra-chromosomal analysis of CDSs' position, $COUSIN_{59}$ and GC3 content in *H. sapiens* are given in Figures 13 (chromosomes 1 to 4), 14 (chromosomes 5 to 8), 15 (chromosomes 9 to 12), 16 (chromosomes 13 to 16), 17 (chromosomes 17 to 20), 18 (chromosomes 21 to Y),

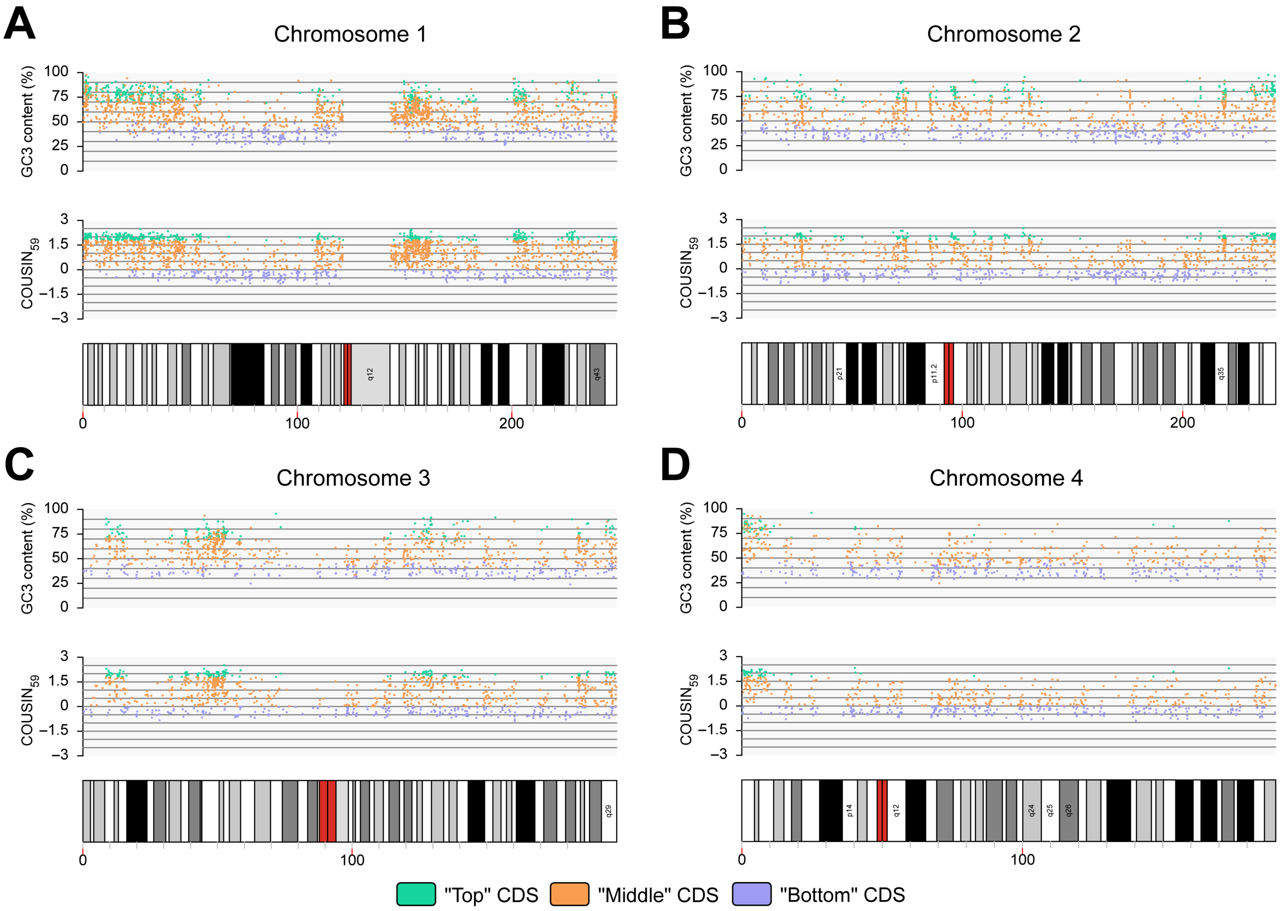


Figure 13: Study of CDSs from *H. sapiens* chromosomes 1 (A), 2 (B), 3 (C) and 4 (D). For each chromosome, GC3 content scores (upper panel), $COUSIN_{59}$ scores (middle panel) and structural information (lower panel) are given. Each dot represents a "Top" (cyan), "Middle" (orange) or "Bottom" (purple) CDSs. The position of CDSs along the related chromosome is given thanks to the x-axis scaled on the graph 1. This scale indicates the distance in megabases from the p-terminal to the q-terminal end of the chromosome. In graphs 3, structural information of chromosomes are centromeres (red), GC-rich and AT-rich isochores (white to black) and chromosomes particularities such as secondary constrictions (blue).

#### 7 Statistics on *G. gallus* and *H. sapiens* by chromosomes

Statistics by chromosomes are given in Tables 3 (*G. gallus*) and 4 *H. sapiens*).

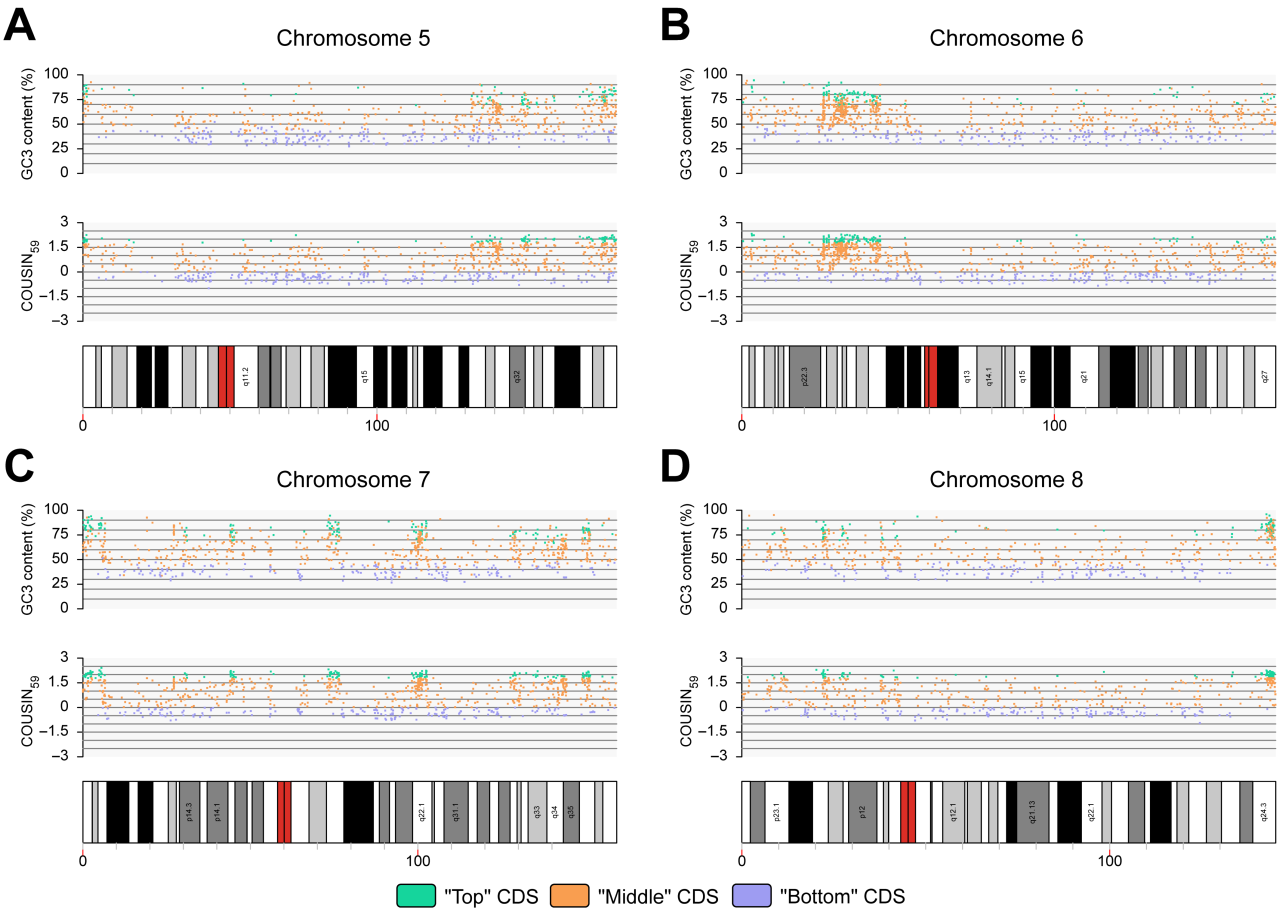


Figure 14: Study of CDSs from *H. sapiens* chromosomes 5 (A), 6 (B), 7 (C) and 8 (D). For each chromosome, GC3 content scores (upper panel), $\text{COUSIN}_{59}$ scores (middle panel) and structural information (lower panel) are given. Each dot represents a "Top" (cyan), "Middle" (orange) or "Bottom" (purple) CDSs. The position of CDSs along the related chromosome is given thanks to the x-axis scaled on the graph 1. This scale indicates the distance in megabases from the p-terminal to the q-terminal end of the chromosome. In graphs 3, structural information of chromosomes are centromeres (red), GC-rich and AT-rich isochores (white to black) and chromosomes particularities such as secondary constrictions (blue).

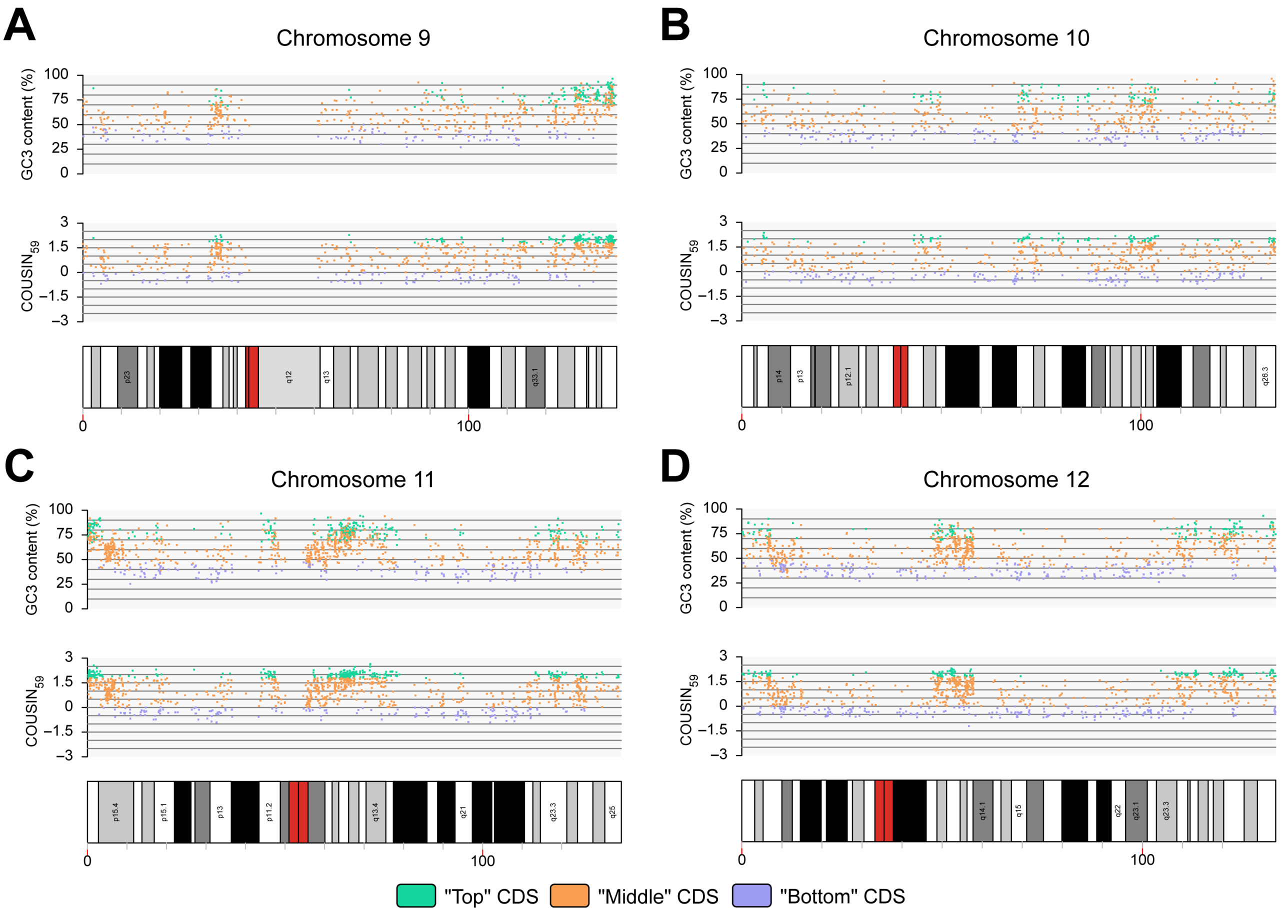


Figure 15: Study of CDSs from *H. sapiens* chromosomes 9 (A), 10 (B), 11 (C) and 12 (D). For each chromosome, GC3 content scores (upper panel), $COUSIN_{59}$ scores (middle panel) and structural information (lower panel) are given. Each dot represents a "Top" (cyan), "Middle" (orange) or "Bottom" (purple) CDSs. The position of CDSs along the related chromosome is given thanks to the x-axis scaled on the graph 1. This scale indicates the distance in megabases from the p-terminal to the q-terminal end of the chromosome. In graphs 3, structural information of chromosomes are centromeres (red), GC-rich and AT-rich isochores (white to black) and chromosomes particularities such as secondary constrictions (blue).

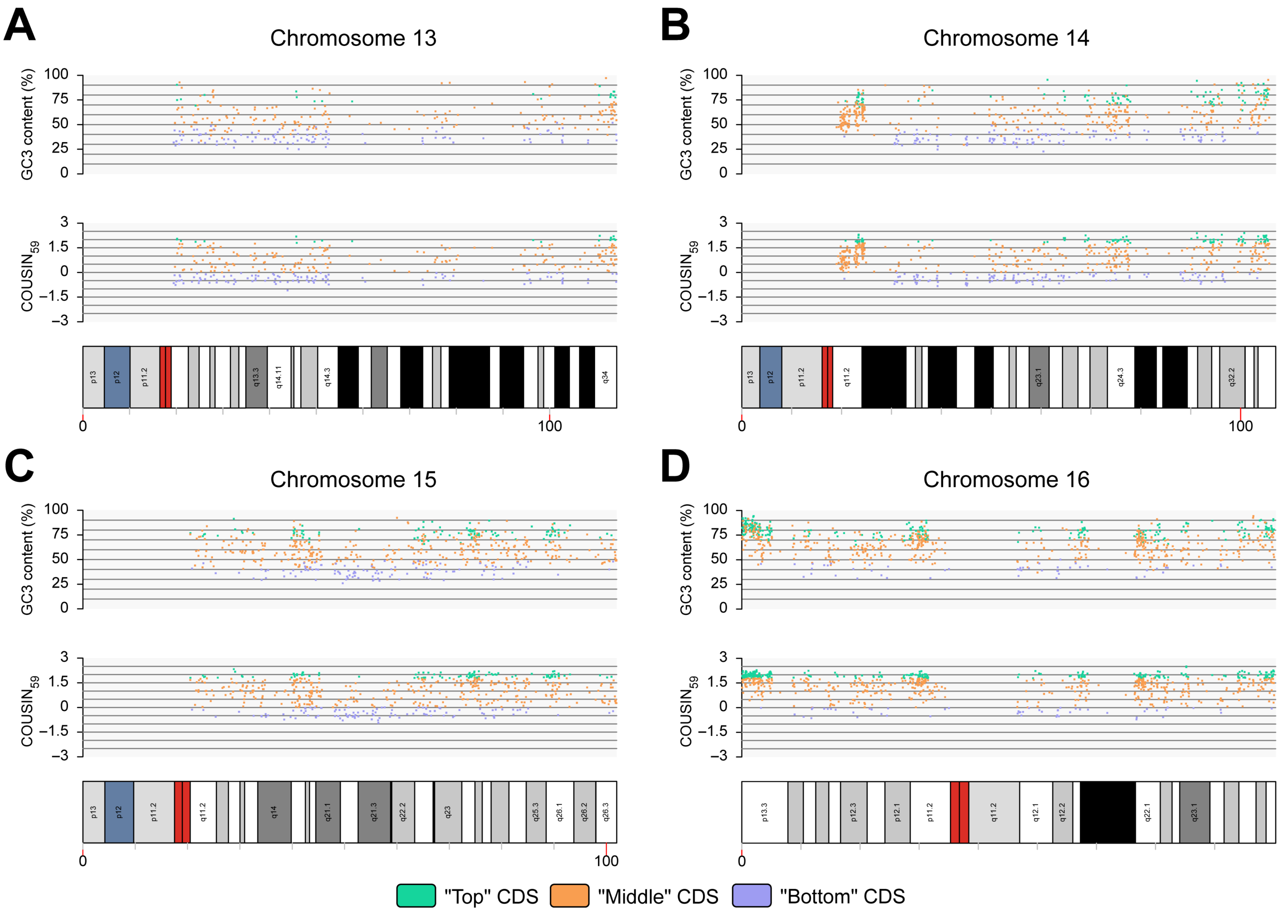


Figure 16: Study of CDSs from *H. sapiens* chromosomes 13 (A), 14 (B), 15 (C) and 16 (D). For each chromosome, GC3 content scores (upper panel), $COUSIN_{59}$ scores (middle panel) and structural information (lower panel) are given. Each dot represents a "Top" (cyan), "Middle" (orange) or "Bottom" (purple) CDSs. The position of CDSs along the related chromosome is given thanks to the x-axis scaled on the graph 1. This scale indicates the distance in megabases from the p-terminal to the q-terminal end of the chromosome. In graphs 3, structural information of chromosomes are centromeres (red), GC-rich and AT-rich isochores (white to black) and chromosomes particularities such as secondary constrictions (blue).

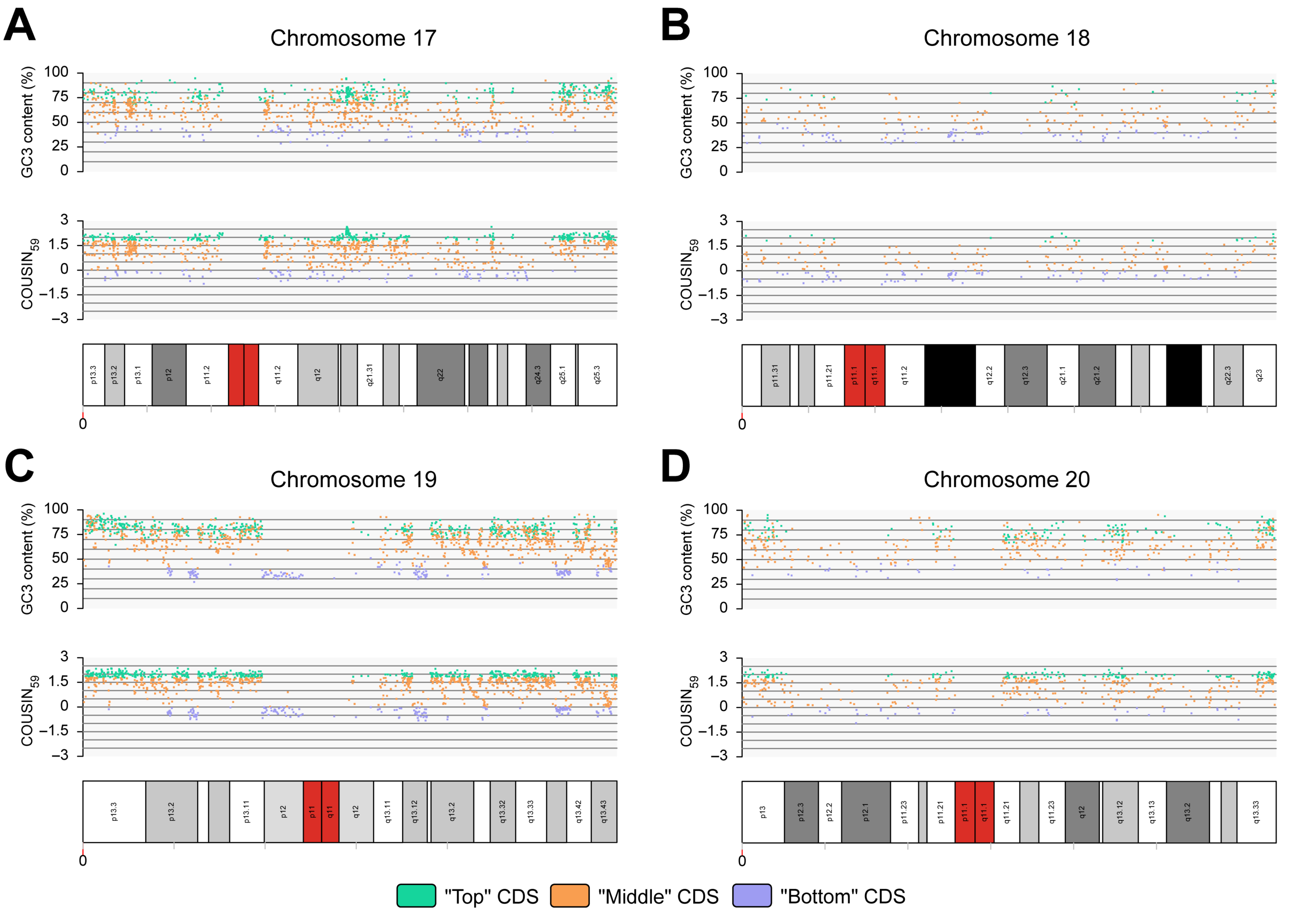


Figure 17: Study of CDSs from *H. sapiens* chromosomes 17 (A), 18 (B), 19 (C) and 20 (D). For each chromosome, GC3 content scores (upper panel), $COUSIN_{59}$ scores (middle panel) and structural information (lower panel) are given. Each dot represents a "Top" (cyan), "Middle" (orange) or "Bottom" (purple) CDSs. The position of CDSs along the related chromosome is given thanks to the x-axis scaled on the graph 1. This scale indicates the distance in megabases from the p-terminal to the q-terminal end of the chromosome. In graphs 3, structural information of chromosomes are centromeres (red), GC-rich and AT-rich isochores (white to black) and chromosomes particularities such as secondary constrictions (blue).

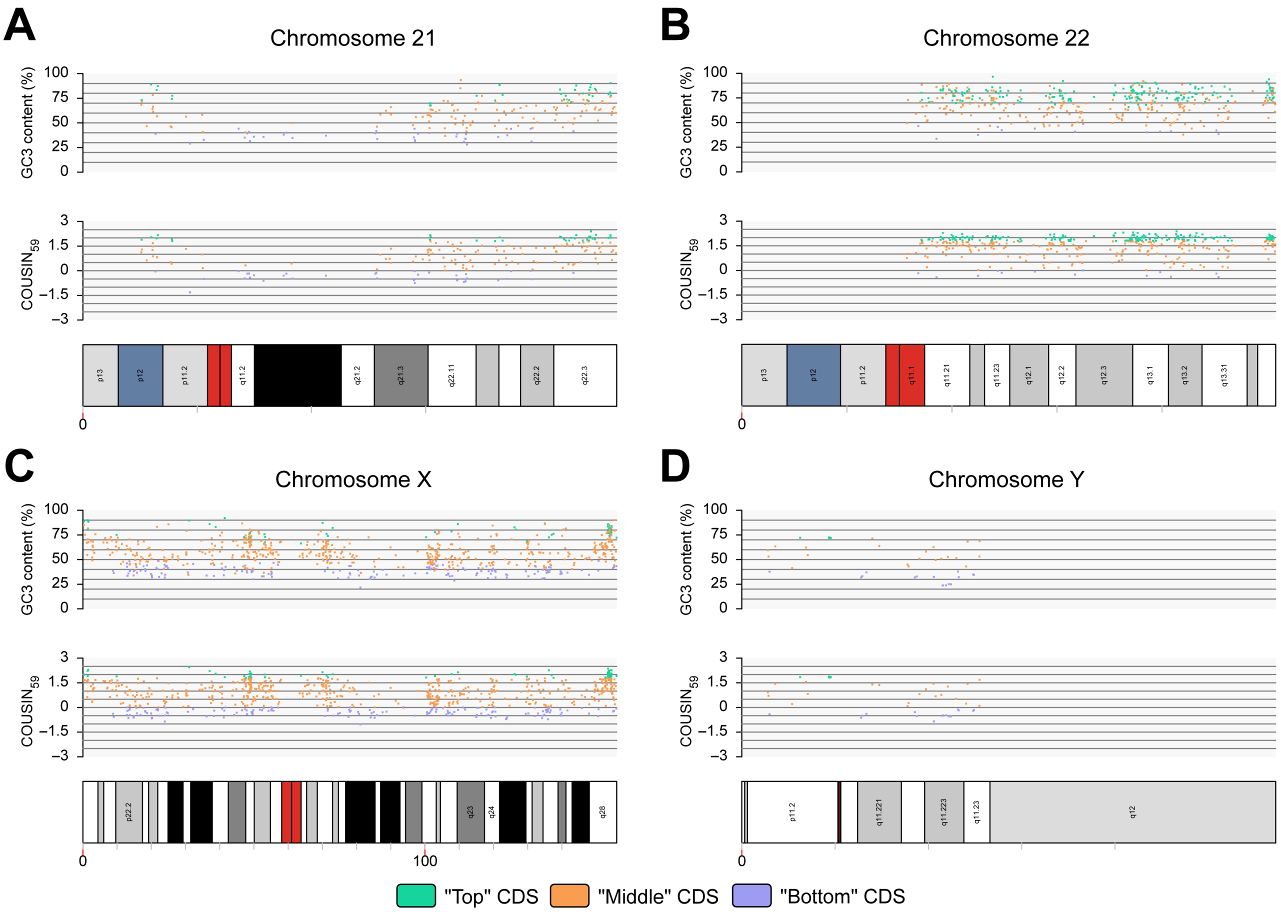


Figure 18: Study of CDSs from *H. sapiens* chromosomes 21 (A), 22 (B), X (C) and Y (D). For each chromosome, GC3 content scores (upper panel), $COUSIN_{59}$ scores (middle panel) and structural information (lower panel) are given. Each dot represents a "Top" (cyan), "Middle" (orange) or "Bottom" (purple) CDSs. The position of CDSs along the related chromosome is given thanks to the x-axis scaled on the graph 1. This scale indicates the distance in megabases from the p-terminal to the q-terminal end of the chromosome. In graphs 3, structural information of chromosomes are centromeres (red), GC-rich and AT-rich isochores (white to black) and chromosomes particularities such as secondary constrictions (blue).

Table 3: Size, number of CDSs, Huber-M estimator values and MAD values for GC3 and $COUSIN_{59}$ among *G. gallus* chromosomes

| Chromosome | Size (Mb) | CDSs | Huber-M estimator (GC3) | MAD (+/-)(GC3) | Huber-M estimator ($COUSIN_{59}$) | MAD (+/-) ($COUSIN_{59}$) |
|---|---|---|---|---|---|---|
| **1** | 197.61 | 2025 | 53.809 | 14.184 | 0.783 | 0.721 |
| **2** | 149.68 | 1315 | 51.390 | 12.507 | 0.650 | 0.581 |
| **3** | 110.84 | 1124 | 53.123 | 13.979 | 0.713 | 0.656 |
| **4** | 91.32 | 1094 | 55.994 | 16.405 | 0.847 | 0.742 |
| **5** | 59.81 | 920 | 58.257 | 17.630 | 0.920 | 0.754 |
| **6** | 36.37 | 520 | 57.453 | 16.860 | 0.865 | 0.690 |
| **7** | 36.74 | 470 | 54.307 | 15.302 | 0.777 | 0.693 |
| **8** | 30.22 | 491 | 57.090 | 19.107 | 0.864 | 0.778 |
| **9** | 24.15 | 415 | 59.799 | 18.996 | 0.962 | 0.752 |
| **10** | 21.12 | 400 | 60.371 | 19.899 | 0.960 | 0.766 |
| **11** | 20.20 | 356 | 63.477 | 22.737 | 1.073 | 0.740 |
| **12** | 20.39 | 340 | 63.982 | 22.446 | 1.040 | 0.685 |
| **13** | 19.17 | 354 | 65.735 | 20.168 | 1.114 | 0.601 |
| **14** | 16.22 | 392 | 63.758 | 18.663 | 1.119 | 0.627 |
| **15** | 13.06 | 347 | 63.616 | 18.173 | 1.110 | 0.603 |
| **16** | 2.84 | 133 | 70.255 | 10.512 | 1.282 | 0.366 |
| **17** | 10.76 | 284 | 66.218 | 16.454 | 1.226 | 0.556 |
| **18** | 11.37 | 306 | 66.297 | 18.971 | 1.238 | 0.552 |
| **19** | 10.32 | 324 | 65.713 | 16.761 | 1.159 | 0.571 |
| **20** | 13.90 | 335 | 64.401 | 18.241 | 1.160 | 0.575 |
| **21** | 6.84 | 236 | 65.035 | 17.188 | 1.223 | 0.481 |
| **22** | 5.46 | 186 | 73.800 | 13.378 | 1.407 | 0.359 |
| **23** | 6.15 | 248 | 69.869 | 17.398 | 1.293 | 0.482 |
| **24** | 6.49 | 184 | 68.106 | 15.044 | 1.357 | 0.429 |
| **25** | 3.98 | 259 | 76.435 | 12.214 | 1.436 | 0.371 |
| **26** | 6.06 | 268 | 70.651 | 14.330 | 1.410 | 0.391 |
| **27** | 8.08 | 296 | 75.11 | 13.051 | 1.471 | 0.361 |
| **28** | 5.12 | 308 | 73.135 | 14.508 | 1.369 | 0.386 |
| **30** | 1.82 | 79 | 79.884 | 6.398 | 1.286 | 0.373 |
| **31** | 6.15 | 213 | 67.833 | 4.399 | 1.304 | 0.382 |
| **32** | 0.73 | 55 | 84.613 | 5.690 | 1.356 | 0.282 |
| **33** | 7.82 | 463 | 73.725 | 11.769 | 1.444 | 0.291 |
| **W** | 6.81 | 37 | 46.625 | 13.277 | 0.448 | 0.503 |
| **Z** | 82.53 | 773 | 52.630 | 15.819 | 0.719 | 0.688 |

Table 4: Size, number of CDSs, Huber-M estimator values and MAD values for GC3 and $COUSIN_{59}$ among *H. sapiens* chromosomes

| Chromosome | Size (Mb) | CDSs | Huber-M estimator (GC3) | MAD (+/-)(GC3) | Huber-M estimator ($COUSIN_{59}$) | MAD (+/-) ($COUSIN_{59}$) |
|---|---|---|---|---|---|---|
| **1** | 248.96 | 1926 | 60.149 | 17.351 | 1.009 | 0.927 |
| **2** | 242.19 | 1171 | 55.914 | 20.690 | 0.715 | 1.208 |
| **3** | 198.30 | 999 | 55.727 | 19.118 | 0.763 | 1.159 |
| **4** | 190.22 | 705 | 48.860 | 13.453 | 0.353 | 0.733 |
| **5** | 181.54 | 763 | 54.795 | 19.816 | 0.670 | 1.203 |
| **6** | 170.81 | 958 | 57.093 | 16.877 | 0.817 | 1.021 |
| **7** | 159.35 | 862 | 59.136 | 20.175 | 0.883 | 1.062 |
| **8** | 145.14 | 631 | 57.114 | 20.067 | 0.754 | 1.190 |
| **9** | 38.40 | 748 | 62.175 | 18.501 | 1.079 | 0.892 |
| **10** | 133.80 | 681 | 57.280 | 18.686 | 0.785 | 1.126 |
| **11** | 135.09 | 1211 | 62.116 | 17.302 | 1.110 | 0.874 |
| **12** | 133.28 | 972 | 56.935 | 19.070 | 0.825 | 1.076 |
| **13** | 114.36 | 306 | 52.032 | 17.210 | 0.486 | 1.020 |
| **14** | 107.04 | 548 | 57.810 | 19.008 | 0.849 | 1.083 |
| **15** | 101.99 | 564 | 58.853 | 17.500 | 0.915 | 1.055 |
| **16** | 90.34 | 792 | 68.980 | 13.521 | 1.465 | 0.520 |
| **17** | 83.26 | 1086 | 67.425 | 14.060 | 1.416 | 0.621 |
| **18** | 80.37 | 252 | 52.147 | 15.972 | 0.512 | 0.903 |
| **19** | 58.62 | 1347 | 71.300 | 11.955 | 1.527 | 0.442 |
| **20** | 64.44 | 490 | 67.107 | 14.670 | 1.365 | 0.571 |
| **21** | 46.71 | 187 | 59.427 | 17.314 | 0.935 | 0.971 |
| **22** | 50.82 | 415 | 71.622 | 11.172 | 1.638 | 0.381 |
| **X** | 156.04 | 751 | 55.853 | 16.324 | 0.791 | 0.995 |
| **Y** | 57.23 | 43 | 48.758 | 23.737 | 0.538 | 1.335 |
